# Unconventional secretion of α-synuclein mediated by palmitoylated DNAJC5 oligomers

**DOI:** 10.1101/2022.01.27.477991

**Authors:** Shenjie Wu, Nancy C. Hernandez Villegas, Daniel W. Sirkis, Iona Thomas-Wright, Richard Wade-Martins, Randy Schekman

## Abstract

Alpha-synuclein (α-syn), a major component of Lewy bodies found in Parkinson’s disease (PD) patients, has been found exported outside of cells and may mediate its toxicity via cell-to-cell transmission. Here, we reconstituted soluble, monomeric α-syn secretion by the expression of DnaJ homolog subfamily C member 5 (DNAJC5) in HEK293T cells. DNAJC5 undergoes palmitoylation and anchors on the membrane. Palmitoylation is essential for DNAJC5-induced α-syn secretion, and the secretion is not limited by substrate size or unfolding. Cytosolic α-syn is actively translocated and sequestered in an endosomal membrane compartment in a DNAJC5-dependent manner. Reduction of α-syn secretion caused by a palmitoylation-deficient mutation in DNAJC5 can be reversed by a membrane-targeting peptide fusion-induced oligomerization of DNAJC5. The secretion of endogenous α-syn mediated by DNAJC5 is also found in a human neuroblastoma cell line, SH-SY5Y, differentiated into neurons in the presence of retinoic acid, and in human induced pluripotent stem cell-derived midbrain dopamine neurons. We propose that DNAJC5 forms a palmitoylated oligomer to accommodate and export α-syn.

## Introduction

Parkinson’s disease, the second most common neurodegenerative disease, is characterized by the deposit of clumps of protein aggregate, lipid and damaged organelles known as Lewy bodies (LBs) (Dauer and Przedborski, 2003; Shahmoradian et al., 2019). One of the main constituents of LB is the presynaptic protein α-syn (Stefanis, 2012). α-syn is encoded by the *SNCA* gene and is highly abundant in neurons. As a small, intrinsically disordered protein containing 140 amino acids (AA), α-syn can be divided into three domains, an amphipathic N-terminal domain where most PD-related mutations are located, including A30P, E46K and A53T, a central hydrophobic region known as the non-amyloid-β component (NAC) which is essential for aggregation, and an acidic C-terminal domain (**Fig 1A**) (Alderson and Markley, 2013). α-syn can undergo a conformational change from a disordered monomer to an oligomer (Burre et al., 2014; Lashuel et al., 2002), which can further polymerize to form insoluble fibrils (Guerrero-Ferreira et al., 2019; Guerrero-Ferreira et al., 2018; Strohaker et al., 2019).

**Figure 1.**
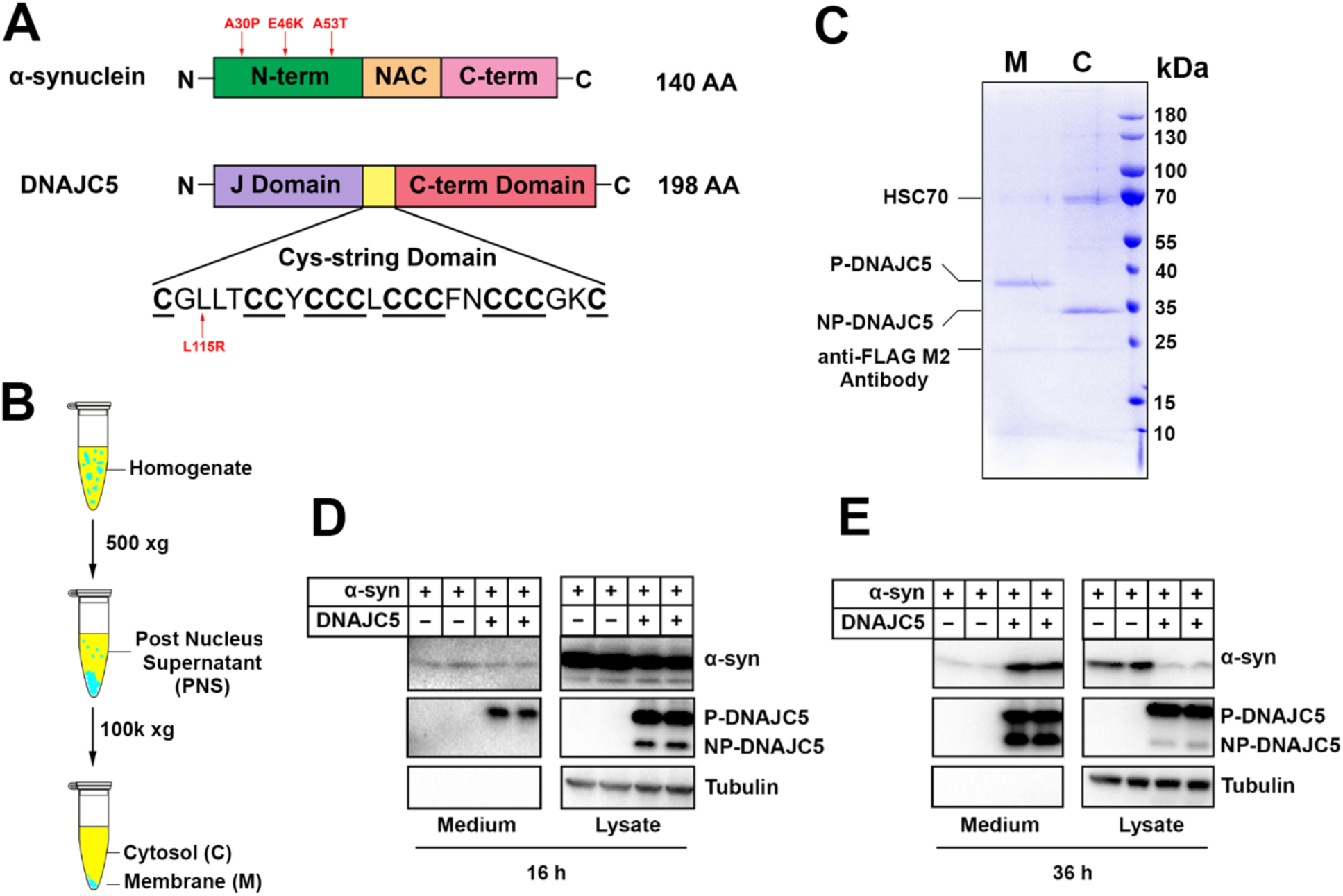
Reconstitution of α-syn secretion regulated by palmitoylated DNAJC5 in HEK293T cells. (A) Schematic diagrams of α-syn and DNAJC5. Domains are highlighted in different colors. Red arrows indicate known disease-causing mutations on each protein. (B) Membrane and cytosol fractionation scheme. Briefly, homogenized HEK293T cells were centrifuged at low speed to prepare a post-nuclear supernatant (PNS). High-speed centrifugation was then performed to separate the sedimentable membrane (M) from cytosol (C). (C) Partition of palmitoylated DNAJC5 (P-DNAJC5) and non-palmitoylated DNAJC5 (NP-DNAJC5) between the membrane (M) and cytosol (C) fractions. DNAJC5 was immunoprecipitated from cytosol and membrane with anti-FLAG resin and evaluated by Coomassie-blue stained SDS-PAGE. (D) α-syn secretion 16 h after transfection. The secretion of P-DNAJC5 in the medium was detected. (E) α-syn secretion 36 h after transfection. NP-DNAJC5 was also secreted in the medium together with α-syn.

In recent years, studies have suggested that α-syn deposits are not static, but rather actively spread during disease progression. Grafted neurons in PD patients developed α-syn positive LBs years after surgery, suggesting host-to-graft pathology propagation (Kordower et al., 2008; Li et al., 2008). Based on analysis of human pathology, the Braak hypothesis posits that α-syn aggregates can spread in a stereotyped manner from the gastrointestinal tract to the brain, causing neuron loss beginning in the brainstem, extending to the midbrain, and finally to the cortex (Braak et al., 2006; Braak et al., 2003). In more recent work, Braak-like transmission of *in vitro* generated α-syn fibrils has been recapitulated in mice and non-human primates (Chu et al., 2019; Chung et al., 2020; Kim et al., 2019; Luk et al., 2012).

Less-well understood is the molecular and cellular basis for the transfer of α-syn between cells. Several mechanisms have been proposed for both monomeric and aggregated α-syn export and transfer, including unconventional exocytosis (Jang et al., 2010; Lee et al., 2005), exosomes (Danzer et al., 2012; Emmanouilidou et al., 2010; Stykel et al., 2021) and membrane nanotubes (Abounit et al., 2016; Scheiblich et al., 2021). The extracellular existence of both monomeric and oligomeric α-syn was confirmed in blood and cerebrospinal fluid (CSF) (Borghi et al., 2000; El-Agnaf et al., 2003). Packaging of different conformational forms of α-syn inside extracellular vesicles (EVs) has been reported but requires further vigorous scrutiny to differentiate membrane vesicles from secreted sedimentable aggregates (Brahic et al., 2016).

DNAJC5, also known as cysteine string protein α (CSPα), is a co-chaperone of HSC70 and has been shown to control the extracellular release of many neurodegenerative disease proteins (Fontaine et al., 2016). This process of unconventional traffic has been termed misfolding-associated protein secretion (MAPS) (Fontaine et al., 2016; Lee et al., 2016) as opposed to conventional secretion initiated by an amino-terminal signal peptide required for secretory and membrane protein translocation into the endoplasmic reticulum (Zhang and Schekman, 2013). DNAJC5 contains three domains – a common N-terminal J-domain conserved among DnaJ proteins, the cysteine-string (CS) central domain which is heavily palmitoylated and anchors the protein to late endosomes, and an overall disordered C-terminal domain (**Fig 1A**). Deletion of DNAJC5 in Drosophila and mice leads to a neurodegenerative phenotype and premature death, indicating that DNAJC5 plays a neuroprotective role in the brain (Zinsmaier, 2010). Transgenic expression of α-syn appears to rescue the neurodegeneration seen on depletion of DNAJC5 (Chandra et al., 2005). A previous study also reported that neuron-derived EVs contain DNAJC5 (Deng et al., 2017). However, the mechanism by which DNAJC5 recognizes and translocates soluble α-syn into a membrane compartment for secretion remains elusive.

In this study, we characterized the mechanism of DNAJC5-induced α-syn secretion in a cell-based secretion assay. Using biochemical characterization and imaging of internalized α-syn in enlarged endosomes as a secretory intermediate, we found previously underappreciated roles of palmitoylation and oligomerization of DNAJC5 in the regulation of α-syn secretion.

## Results

### Reconstitution of DNAJC5-induced α-syn secretion

Previous studies have shown that α-syn secretion can be stimulated by over-expressing DNAJC5 (Fontaine et al., 2016). The CS domain of DNAJC5 plays a role in promoting stable membrane attachment based on its overall hydrophobicity and by enabling post-translational palmitoylation catalyzed by membrane-bound Asp-His-His-Cys (DHHC) family palmitoyltransferases (Greaves and Chamberlain, 2006). Using a common human cell line, HEK293T, we first tested the expression and subcellular localization of DNAJC5 (**Fig 1B**). Further subcellular enrichment of DNAJC5 was characterized using a C-terminal FLAG tag. Coomassie blue staining revealed two bands of low and high mobility on SDS-PAGE in the membrane and cytosolic fractions, respectively (**Fig 1C**). Similar migration profiles of pamitoylated (P-) and non-palmitoylated (NP-) DNAJC5 have been reported (Greaves et al., 2012). We also transfected and fractionated DNAJC5 in other common cell lines including MDA-MB-231 and Hela cells. Compared to DNAJC5 in HEK293T cells, DNAJC5 in MDA-MB-231 and Hela cells appeared predominantly to be in a palmitoylated and membrane-associated form (**Fig 1 supplement 1A**). The mobility of P-DNAJC5 in the membrane fraction shifted to that of NP-DNAJC5 after an overnight depalmitoylation reaction with hydroxylamine (HA) (**Fig 1 supplement 1B**). Thus, we confirm that membrane anchoring of DNAJC5 requires palmitoylation in our assay.

We next co-expressed DNAJC5 together with α-syn to examine their secretion over time. At 16 h after transfection, we detected similar basal-levels secretion of α-syn in both DNAJC5-negative and -positive conditions. Two bands of DNAJC5 corresponding to P-DNAJC5 and NP-DNAJC5 were seen in the lysate, but only the P-DNAJC5 was secreted into the medium (**Fig 1D**). The stimulation of α-syn secretion by DNAJC5 became obvious at a longer incubation time (36 h), and at this time point NP-DNAJC5 was also enriched in the medium (**Fig 1E**). The release of α-syn was not caused by cell death as little to undetected levels of cytoplasmic tubulin found in the culture medium fraction (**Fig 1D&1E**). Cell viability was not affected by transfection of different constructs, as shown by trypan blue staining (**Fig 1 supplement 2A**). In addition to wild-type (WT) α-syn, secretion of several PD-causing α-syn mutant proteins (A30P, E46K and A53T) was also induced to differing levels by expression of DNAJC5 (**Fig 1 supplement 2B-2D**).

In addition to the stimulated secretion of α-syn produced by the expression of exogenous DNAJC5, we examined the dependence of a basal secretion of α-syn on endogenous DNAJC5. We fused α-syn with an N-terminal nanoluciferase (Nluc) (England et al., 2016) for sensitive, quantitative detection (**Fig 1 supplement 3A**). Stimulated secretion of Nluc-α-syn by overexpression of DNAJC5 was confirmed by immunoblot, indicating that Nluc-fusion did not impede α-syn secretion (**Fig 1 supplement 3B**). Without over-expression of DNAJC5, we observed accumulation of Nluc-α-syn signal in the medium over time (**Fig 1 supplement 3C**).

Quercetin is a plant-derived flavonoid that has previously been shown to inhibit DNAJC5-mediated trafficking of a bacterial toxin (**Fig 1 supplement 3D**) (Deruelle et al., 2021). We found that quercetin also inhibited Nluc-α-syn secretion in a dose-dependent manner (**Fig 1 supplement 3E**), implying a role for endogenous DNAJC5 in α-syn secretion. To exclude the off-target effect of quercetin, we created a DNAJC5 CRISPR knockout (KO) cell line (**Fig 1 supplement 3F**). Balfilomycin A1 (BaFA1), a lysosomal ATPase inhibitor, has been shown to stimulate α-syn secretion (**Fig 1 supplement 3G**) (Buratta et al., 2020; Fernandes et al., 2016). BafA1 is also known to stimulate the fusion of lysosomes at the cell surface with the secretion of lysosomal content (Tapper and Sundler, 1995). BaFA1-stimulated α-syn secretion was confirmed in WT HEK293T cells but not observed in DNAJC5 KO cells (**Fig 1 supplement 3H**). Our results suggest that α-syn secretion depends on the basal level of DNAJC5 and is enhanced by overexpression of DNAJC5.

### Characterization of extracellular DNAJC5 and α-syn

In our established assay, DNAJC5 and α-syn co-secrete into the medium (**Fig 1E**). Secreted α-syn has been reported to be encapsulated inside EVs (Danzer et al., 2012). We sought to assess the EV association of secreted α-syn using a medium fractionation protocol based on EV preparations developed by our lab (**Fig 2A**) (Shurtleff et al., 2016). After serial differential centrifugation, α-syn and NP-DNAJC5 remained soluble. In comparison, P-DNAJC5 co-sedimented with other EV markers after 100k xg centrifugation (**Fig 2B**). In addition to WT α-syn, we conducted medium fractionation with several PD-causing α-syn mutants and found they all remained soluble in culture supernatant fractions (**Fig 2 supplement 1A**). To test the EV association of P-DNAJC5, we performed a further sucrose step gradient flotation (**Fig 2 supplement 2A**) with the 100k high-speed pellet fraction (**Fig 2 supplement 2B**). P-DNAJC5 equilibrated with other EV markers to the 10%/40% interface expected for buoyant EVs (**Fig 2 supplement 2B**). We conclude that the secreted α-syn induced by DNAJC5 is neither membrane bound nor in a sedimentable fibrillar form.

**Figure 2.**
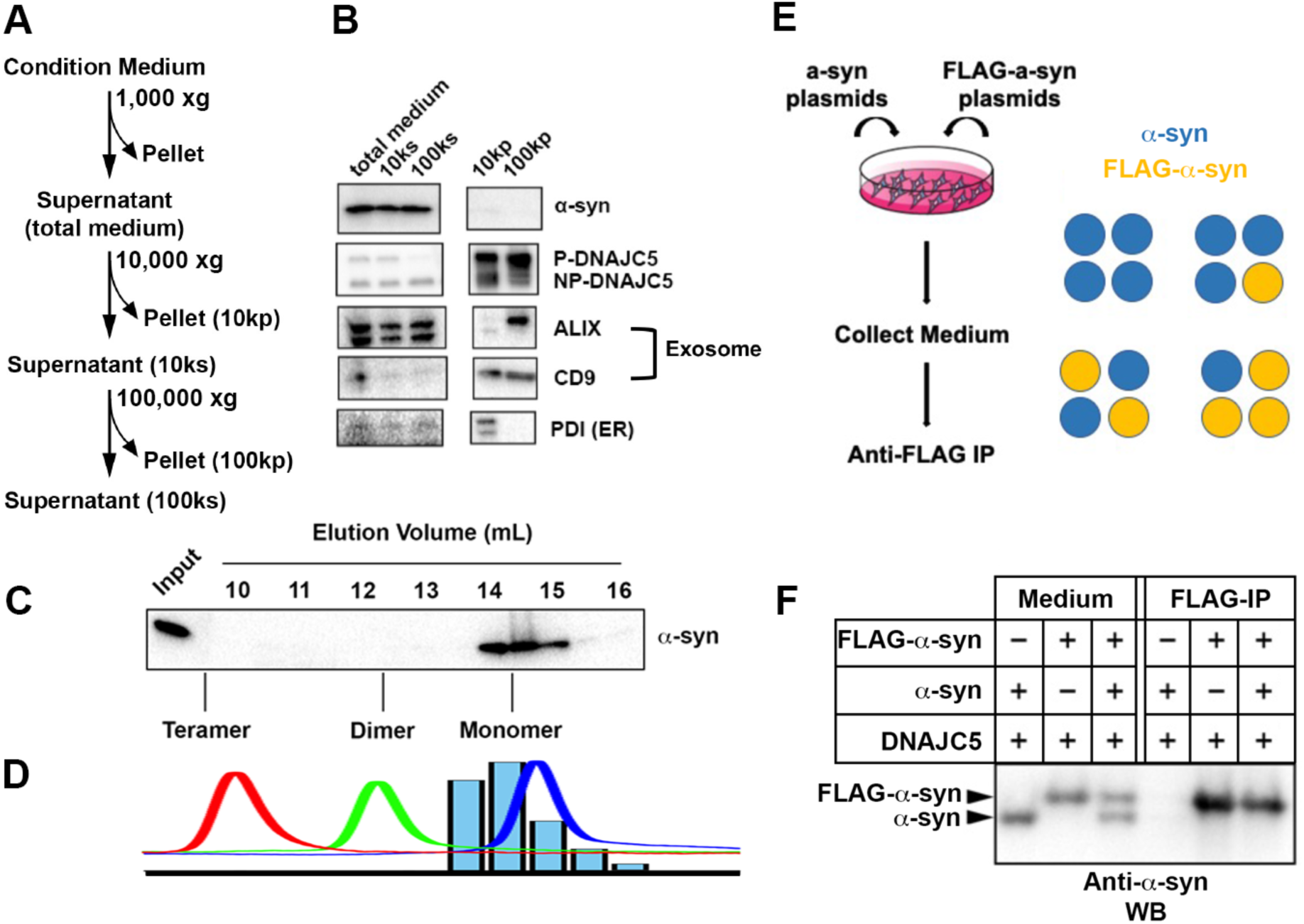
Characterization of secreted α-syn. (A) Medium fractionation scheme. (B) Secreted α-syn was soluble. Differential centrifugation was performed with conditioned medium from HEK293T cells transfected with DNAJC5 and α-syn. Alix and CD9, exosome markers. PDI, an endoplasmic reticulum (ER) marker, was used as exosome-negative control. (C) Gel filtration fractionation of medium. Conditioned medium was concentrated and subjected to gel filtration fractionation. Fractions were evaluated by anti-α-syn immunoblot. (D) Chromatograms of tandem α-syn monomer (blue curve), dimer (green curve) and tetramer (red curve) were overlaid. In comparison, the relative intensity of secreted α-syn in each fraction was plotted as blue bars. (E) Schematic diagram of co-immunoprecipitation (co-IP) of secreted α-syn and FLAG-α-syn. Shown here is possible interaction between α-syn (blue circle) and FLAG-α-syn (yellow circle) in a representative tetrameric conformation. (F) Anti-FLAG immunoprecipitation (FLAG-IP) of media from cells transfected with indicated plasmids. Both the medium input and FLAG-IP samples were evaluated with anti-α-syn immunoblot (anti-α-syn WB).

Next we sought to characterize the conformation of secreted soluble α-syn. The medium containing secreted α-syn was pooled, concentrated and applied to a gel filtration column. Extracellular α-syn eluted from the column at around 60% of the column volume (**Fig 2C**), similar to the elution volume of purified monomeric α-syn (**Fig 2D and Fig 2 supplement 3E&F**). By an orthogonal assay, we examined the interaction between tagged and untagged forms of secreted α-syn as an indicator of oligomerization. In this assay, equal amounts of plasmids expressing FLAG-tagged α-syn and non-tagged α-syn were co-transfected in HEK293T cells and the medium was collected and incubated with anti-FLAG M2 beads for to detect co-immunoprecipitation of the two forms (IP) (**Fig 2E**). FLAG-tagged α-syn migrated more slowly than non-tagged α-syn as detected in samples of the culture medium (**Fig 2F**). We found that only the FLAG-tagged α-syn was immuno-precipitated (**Fig 2F**), suggesting no stable interaction between the two species. The gel filtration and IP assays reinforced our conclusion that α-syn is secreted as a monomer.

### Secretion of α-syn requires palmitoylation of DNAJC5

Membrane targeting of DNAJC5 is dependent upon palmitoylation (Greaves et al., 2008). Two specific mutations, L115R and L116Δ in the CS domain, cause adult-onset neuronal ceroid lipofuscinosis (NCL), a type of neurodegenerative disorder (Benitez et al., 2011). NCL mutations reduce the level of DNAJC5 palmitoylation and promote aggregation of the protein (Diez-Ardanuy et al., 2017). We perturbed DNAJC5 palmitoylation by either introducing the palmitoylation-deficient mutation L115R or treating cells with the competitive palmitoyl transferase inhibitor, 2-bromopalmitic acid (2-BA) (Resh, 2006), and subsequently examined the influence on DNAJC5 membrane association. DNAJC5 palmitoylation largely decreased in the L115R mutant or upon 2-BA treatment. Correspondingly, NP-DNAJC5 accumulated in the cytosol (**Fig 3A&3B**).

**Figure 3.**
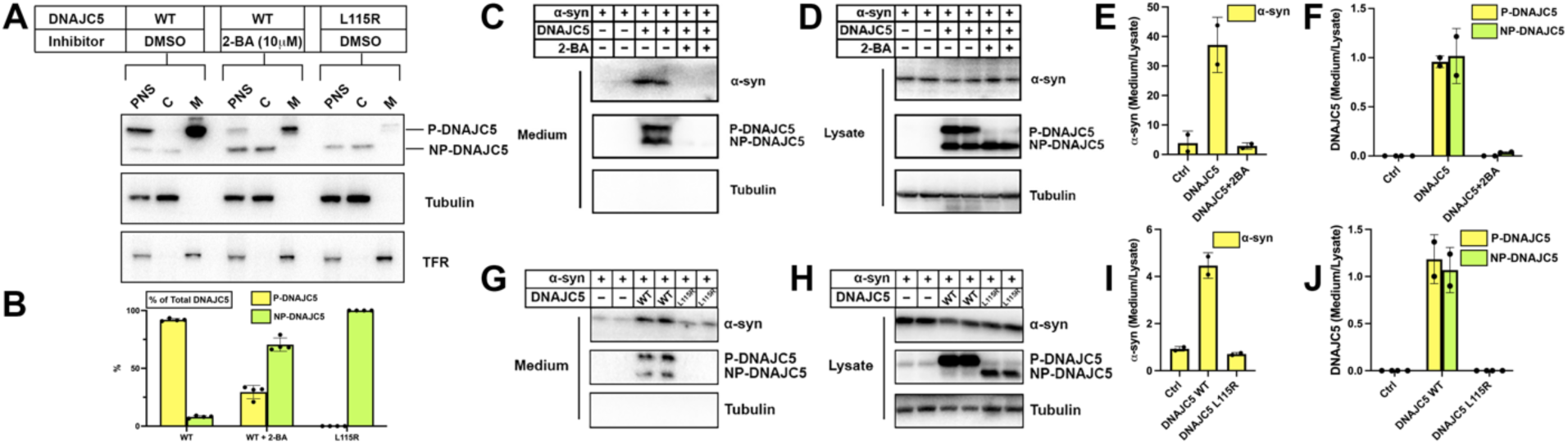
Disruption of palmitoylation of DNAJC5 inhibited α-syn secretion. (A) Inhibition of DNAJC5 palmitoylation by 2-bromopalmitic acid (2-BA) or introduced mutation L115R. Cellular fractionation was performed with HEK293T cells transfected with WT DNAJC5 and treated with 10 μm 2-BA, or transfected with DNAJC5 L115R mutant. TFR, transferrin receptor. PNS, post-nuclear supernatant. C, cytosol. M, membrane. (B) Quantification of the percentage of P-DNAJC5 and NP-DNAJC5 in different conditions as shown in (A). Error bars represent standard deviations of 3 experiments. (C) α-syn secretion was blocked with 2-BA treatment. HEK293T cells transfected with indicated plasmids were treated with DMSO or 10 μm 2-BA. Media fractions were collected and secretion was evaluated by SDS-PAGE and immunoblot. (D) Palmitoylation of DNAJC5 was blocked in HEK293T cells treated with 2-BA. (E) Quantification of normalized α-syn secretion in HEK293T cells after 2-BA treatment. The quantification was based on immunoblot in (C) and (D). The α-syn secretion was calculated as the amount of α-syn in media divided by the amount in lysate. (F) Quantification of normalized DNAJC5 secretion in HEK293T cells after 2-BA treatment. The quantification was based on immunoblot in (C) and (D). The DNAJC5 secretion was calculated as the amount of DNAJC5 in media divided by the amount in lysate. (G) DNAJC5 L115R mutant reduced α-syn secretion compared with WT DNAJC5. Secretion assay with HEK293T cells transfected with indicated plasmids encoding DNAJC5 variant was performed similar to (C). (H) DNAJC5 L115R was non-palmitoylated in HEK293T cells. (I) Quantification of normalized α-syn secretion in HEK293T cells transfected with DNAJC5 L115R mutant. The quantification was based on immunoblot in (G) and (H). (J) Quantification of normalized DNAJC5 secretion in HEK293T cells transfected with DNAJC5 L115R mutant. The quantification was based on immunoblot in (G) and (H).

Having confirmed palmitoylation inhibition by 2-BA and the L115R mutation, we next examined their effect on α-syn secretion. Upon 10 μM 2-BA treatment, α-syn and DNAJC5 secretion were abolished (**Fig 3C, 3E&3F**). The efficacy of the inhibitor was validated by the disappearance of the low-mobility band corresponding to P-DNAJC5 in the cell lysate (**Fig 3D**). Furthermore, 2-BA inhibited DNAJC5 palmitoylation (**Fig 3 supplement 1C&1E**) and α-syn secretion in a concentration-dependent manner (**Fig 3 supplement 1B&1D**). Likewise, α-syn secretion was reduced by the palmitoylation-deficient DNAJC5 (L115R) compared with DNAJC5 (WT) (**Fig 3G-3J**). DNAJC5 has been proposed to function downstream of the deubiquitinase USP19 in the MAPS pathway (Xu et al., 2018). In agreement with the model, DNAJC5 carrying either NCL mutation L115R or L116Δ had no palmitoylated protein band detected in the lysate (**Fig 3 supplement 2B**) and blocked USP19-stimulated α-syn secretion (**Fig 3 supplement 2A&2C**). These results establish that palmitoylation is essential for DNAJC5 membrane association and function in stimulating α-syn secretion.

A previous study reported an altered distribution of DNAJC5 mutant protein to the Golgi apparatus and cytosol (Noskova et al., 2011). In confocal immunofluorescence (IF) images, we confirmed that WT DNAJC5 did not colocalize with the Golgi marker GM130 whereas both the L115R mutant and 2-BA treated WT cells partially retained DNAJC5 in puncta coincident with the Golgi marker GM130 (**Fig 3 supplement 3A,B&C**). We conclude that α-syn secretion depends upon an appropriate subcellular organelle localization of DNAJC5.

### DNAJC5-dependent internalization of α-syn into enlarged endosomes

DNAJC5 has been reported under normal conditions to be associated with late endosomes (Lee et al., 2018). Using confocal IF, we found colocalization between endogenous DNAJC5 and the late-endosomal marker CD63 (**Fig 4 supplement 1A**). To visualize the topological localization of DNAJC5 and α-syn inside or outside endosomes, we turned to a U2OS cell line expressing a fluorescent protein-fused, constitutively active form of Rab5 (Rab5^Q79L^) (Bohdanowicz et al., 2012). As a positive control, CD63 localized to the lumen of enlarged endosomes labeled by mCherry-Rab5^Q79L^(**Fig 4 supplement 1B**). We labeled DNAJC5 with the self-labeling HaloTag for multiple choices of color in live cell imaging (Los et al., 2008). Both diffuse and punctate DNAJC5 localized to the lumen of enlarged endosomes (**Fig 4A**). Unlike WT DNAJC5, the DNAJC5 L115R mutant became disperse in the cytosol, rather than being internalized into enlarged endosomes (**Fig 4 supplement 2A**). In contrast, mNeonGreen (mNG)-fused α-syn showed diffuse localization in both the cytosol and nucleus but was completely excluded from enlarged endosomes in L115R mutant cells (**Fig 4B**). Notably, we observed the entry of α-syn into enlarged endosomes containing internalized DNAJC5, implying the translocation of α-syn into the membrane compartment required DNAJC5 (**Fig 4C**). The ratio of α-syn-containing endosomes in cells increased significantly with DNAJC5 overexpression (**Fig 4H)**. With no luminal localization inside enlarged endosomes, the DNAJC5 L115R mutants also failed to induce entry of α-syn into the same compartments (**Fig 4 supplement 2B**). As an independent test of the localization suggested by the imaging results, we applied cell fractionation to separate membranes of DNAJC5- and α-syn-expressing cells (**Fig 4D**). Both DNAJC5 and α-syn were enriched in a 25k membrane pellet fraction (**Fig 4E**). In a protease protection assay with 25k sedimented membranes, we found that α-syn and DNAJC5 were partially resistant to digestion by proteinase K in the absence but not in the presence of Triton X-100 consistent with the conclusion that about half of the proteins were sequestered within membrane compartments (**Fig 4F&4G)**. Our visual inspection and quantification results are consistent with a membrane translocation role for DNAJC5 prior to α-syn secretion.

**Figure 4.**
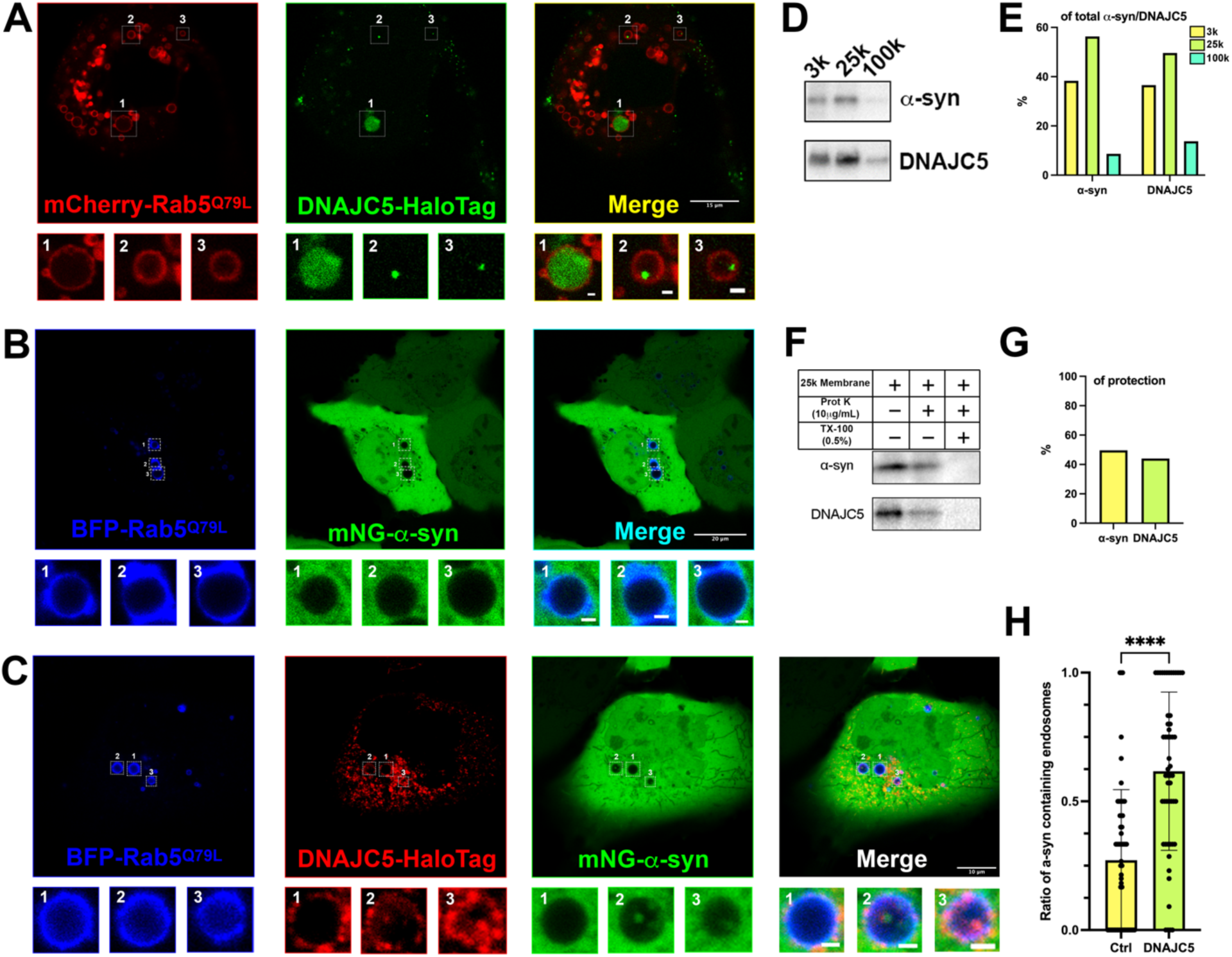
Topological localization of α-syn and DNAJC5 in enlarged endosomes. (A) DNAJC5 was internalized inside enlarged endosomes. Live U2OS cells expressing mCherry-Rab5^Q79L^(red) showed circular enlarged endosomes labeled by Rab5 mutant. DNAJC5-HaloTag (green) was visualized by addition of HaloTag^®^ Oregon Green^®^ Ligand. Representative enlarged endosomes show diffuse (1) or punctate (2 and 3) internalized DNAJC5. Scale bar: 15 μm in overviews and 1 μm in magnified insets. (B) α-syn was excluded from enlarged endosomes. In live U2OS cells, expression of BFP-Rab5^Q79L^ (blue) produced enlarged endosomes of similar morphology compared with mCherry-Rab5^Q79L^. mNeonGreen-α-syn (mNG-α-syn, green) was expressed both in the nucleus and cytosol. No mNG-α-syn was found inside enlarged endosomes (1, 2, 3). Scale bar: 20 μm in overviews and 1 μm in magnified insets. (C) α-syn enters into enlarged endosomes in the presence of DNAJC5. DNAJC5-HaloTag (red) and mNG-α-syn (green) were co-expressed in U2OS cells carrying BFP-Rab5^Q79L^ (blue) mutant and imaged. No mNG-α-syn was internalized in endosome without DNAJC5-HaloTag inside (1). In contrast, mNG-α-syn was found inside endosomes with DNAJC5-HaloTag inside (2 and 3). Scale bar: 10 μm in overviews and 1 μm in magnified insets. (D) α-syn and DNAJC5 co-sedimented in membrane fractionation. HEK293T cell homogenate was sequentially centrifuged at increasing velocity from 3000g (3k), 25,000g (25k) and 100,000g (100k). The 25k membrane fraction had the highest amount of both α-syn and DNAJC5. (E) Quantification of the membrane fractionation results in (D). (F) Proteinase K protection assay of 25k membrane containing α-syn and DNAJC5. (G) Quantification of the proteinase K protection assay in (F). (H) Quantification of the ratio of α-syn containing endosomes in control cells (no-DNAJC5 transfection) or cells co-transfected with DNAJC5. More than 100 enlarged endosomes were counted in each group. Error bars represent standard deviations. ****p value<0.0001, two-tailed t test.

### Size and unfolding are dispensable for α-syn secretion

In our medium fractionation assay, secreted α-syn in the extracellular space was characterized as a soluble monomer (**Fig 2**). We generated a series of tandem repeats of α-syn to mimic its oligomeric states (**Fig 5A**) (Dong et al., 2018). On SDS-PAGE, α-syn tandem repeats showed a larger apparent size than their predicted molecular weights, possibly caused by their extended conformation as intrinsically disordered proteins (**Fig 2 supplement 2**). We first determined that these α-syn tandem repeats could also be secreted upon overexpression of DNAJC5 (**Fig 5B**), indicating that DNAJC5 can accommodate α-syn of different sizes. Fractionation of the growth medium showed that secreted tandem repeats were also soluble (**Fig 5 supplement 1**).

**Figure 5.**
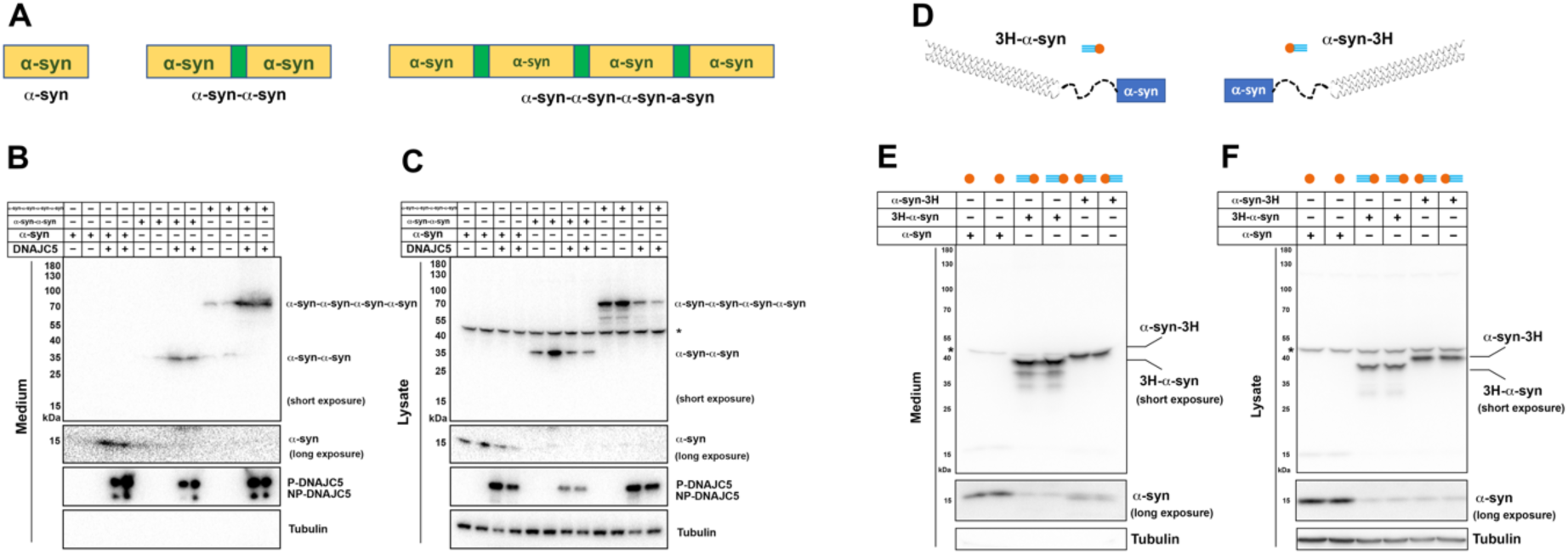
Secretion of tandem α-syn oligomers and α-syn fused with thermostable helix bundle protein. (A) Schematic diagrams of tandem α-syn oligomers. α-syn protomers (yellow) were linked head to tail by flexible linker (green) to mimic increased size of α-syn oligomers. (B) Secretion of tandem α-syn oligomers in medium. Secretion assay was performed with media from HEK293T cells transfected with indicated tandem α-syn oligomers. Tandem α-syn oligomers are more sensitively detected by immunoblot which were exposed for shorter time compared with WT α-syn. (C) Expression of tandem α-syn oligomers in HEK293T cells. *a non-specific band. (D) Schematic diagrams of N-terminal fused and C-terminal fused thermostable three helix bundle (3H-) α-syn. 3H shown as three blue dashes, α-syn shown as orange circle. (E) Secretion of 3H-α-syn and α-syn-3H in medium. Secretion assay was performed with media from HEK293T cells transfected with indicated 3H-fused α-syn constructs. *a non-specific band. (F) Expression of 3H-α-syn and α-syn-3H in HEK293T cells. *a non-specific band.

In conventional and many unconventional secretion processes, the secreted proteins undergo unfolding prior to translocation through a narrow channel across the hydrophobic membrane barrier (Rapoport et al., 2017). Recent progress in protein design has allowed the synthesis of super-folded protein constructs (Kuhlman and Bradley, 2019). For example, a three helix-bundle protein (3H) designed in the lowest-energy arrangements displayed extreme thermodynamic stability and remained folded even in non-physiological denaturing conditions (Huang et al., 2014). Given that mitochondrial protein import is dependent on protein unfolding (Neupert, 1997), we tested the effect of 3H insertion on the import of mitochondrial matrix enzyme ornithine transcarbamylase (OTC) (Horwich et al., 1985; Yano et al., 1997). As a control, we created a pOTC leader peptide fused to GFP (**Fig 5 supplement 2A**). pOTC-GFP was enriched in a mitochondria-containing particulate (P) fraction compared to non-tagged GFP. However, a construct in which 3H was inserted between the leader sequence and GFP resulted in 80% of the fusion protein retained in the soluble fraction (**Fig 5 supplement 2B&2C**). We then used proteinase K protection to assess the topology of pOTC-3H-GFP associated with crude mitochondria (**Fig 5 supplement 2D**). About 80% of citrate synthase (CS), a known mitochondria matrix protein, was protected from proteinase K. In contrast, neither the mitochondrial outer membrane protein Tom20 nor pOTC-3H-GFP was protected, suggesting 3H prevented the translocation of GFP into mitochondria (**Fig 5 supplement 2E&2F**). Using a similar approach, we fused 3H to either the N- or C-terminus of α-syn to impede the unfolding process (**Fig 5D**). These α-syn fusion proteins were expressed and secreted normally into the growth medium (**Fig 5E&F**). These data suggest that protein unfolding is dispensable for α-syn secretion.

### XPACK fusion rescues DNAJC5 L115R secretion deficiency by induced oligomerization

DNAJC5 has been reported to have an intrinsic propensity to form SDS-resistant oligomers (Zhang and Chandra, 2014). In a whole gel immunoblot of extracellular DNAJC5, we noticed many diffuse, ladder-like bands that migrated more slowly than the two corresponding to P-DNAJC5 and NP-DNAJC5 (**Fig 6A**), possibly higher molecular weight (HMW) oligomers. This apparent oligomerization of DNAJC5 became more obvious when the J domain was deleted (**Fig 6 supplement 1A**). The migration of HMW-DNAJC5 was not altered in samples heated in the presence of a reducing agent (**Fig 6 supplement 1B**). To assess the size of these HMW species of DNAJC5, we evaluated a cell lysate by gel filtration chromatography. HMW-DNAJC5 fractionated according to its apparent size, forming a stair-like pattern on the DNAJC5 immunoblot (**Fig 6B**). HMW-DNAJC5 chromatographed within the gel filtration column volume, consistent with discrete protein species rather than aggregates (**Fig 6B**). These results suggest the presence of higher-order, SDS-resistant, non-disulfide-bonded DNAJC5 oligomers both in intracellular and extracellular fractions.

**Figure 6.**
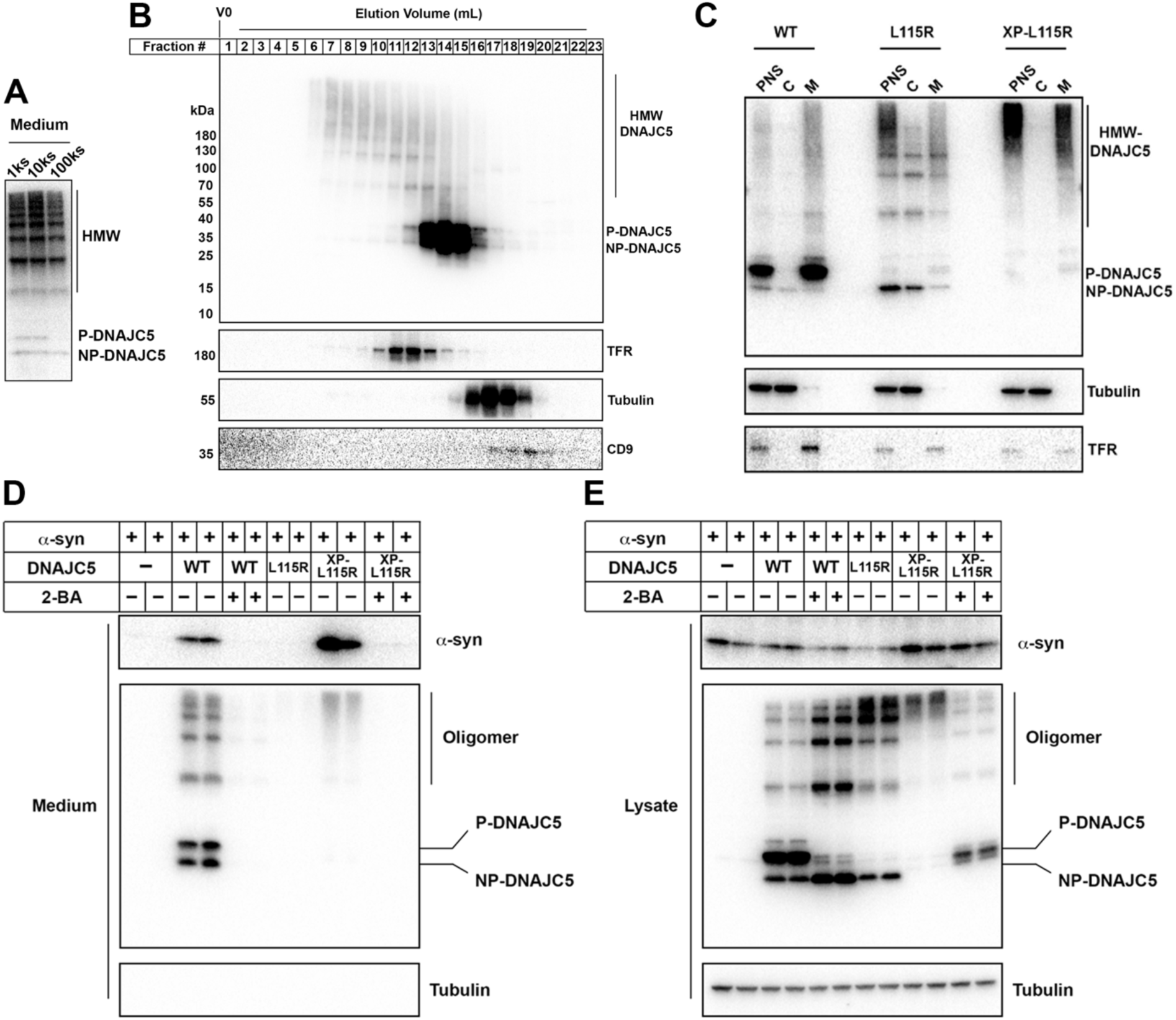
XPACK (XP)-induced DNAJC5 L115R oligomerization rescued α-syn secretion. (A) Ladder pattern of higher molecular weight (HMW) DNAJC5 oligomers in the medium. Medium from HEK293T cell culture transfected with DNAJC5 was centrifuged at 1000 (1k) g, 10,000 (10k) g and 100,000 (100k) g, followed by SDS-PAGE and immunoblot of supernatant (s) fractions at each centrifugation step. (B) Fractionation of HMW-DNAJC5 with gel filtration. HEK293T cells transfected with DNAJC5 were lysed, clarified and subjected to gel filtration. HMW-DNAJC5 of different sizes were separated based on their corresponding molecular weight. (C) XP-DNAJC5 L115R mutant forms a membrane-bound oligomer. Cellular fractionation was performed with HEK293T cells transfected with indicated DNAJC5 variants. Note the substantial change of electrophoretic mobility of XP-DNAJC5 L115R on SDS-PAGE. (D) α-syn secretion induced by XP-DNAJC5 L115R. Secretion assay was performed with HEK293T cells transfected with indicated plasmids. Ten μm 2-BA was used to block induced α-syn. (E) Expression of α-syn and DNAJC5 variants in HEK293T cells. Note the substantial change in electrophoretic mobility of 2-BA-treated XP-DNAJC5 L115R on SDS-PAGE.

XPACK (XP) is a membrane-targeting peptide sequence used widely in studies of cargo loading into exosomes and delivery to target cells of choice (Yim et al., 2016). The exosome loading process by XP is also dependent on two lipidation reactions – myristoylation on the first glycine and palmitoylation on the second cysteine (**Fig 6 supplement 1C**) (Zacharias et al., 2002). Given the similarity of membrane localization and lipidation between XP and the CS domain of DNAJC5, we examined α-syn secretion in cells expressing an XP-DNAJC5 fusion. An N-terminal XP fusion (XP-WT) resulted in the expression of a species that migrated at the position of P-DNAJC5, in contrast to the two species representing P- and NP-DNAJC5 in the WT DNAJC5 sample (**Fig 6 supplement 1D**). Again in contrast to WT DNAJC5, XP-DNAJC5 was exclusively associated with the sedimentable membrane fraction (**Fig 6 supplement 1E**). This suggested that XP-mediated lipidation was highly efficient and possibly irreversible. Formation of the lower electrophoretic mobility and membrane associated form of XP-DNAJC5 was blocked by 2-BA treatment, indicating XP lipidation included palmitoylation (**Fig 6 supplement 1E**). XP fusion to the palmitoylation-deficient mutant of DNAJC5 (L115R) did not produce a species that migrated at the position of P-DNAJC5 but resulted in several less abundant species that migrated between the positions of NP- and P-DNAJC5 (**Fig 6 supplement 1D**). We introduced a serine to leucine point mutation in the XPACK sequence which was predicted to block lipidation (dead XPACK, DXP) (**Fig 6 supplement 1C**). The DXP-DNAJC5 L115R species had the same mobility as NP-DNAJC5 (**Fig 6 supplement 1D**).

We conducted cellular fractionation on lysates of cells expressing DNAJC5 XP-L115R. XP-L115R was highly enriched in the membrane fraction, likely as a result of XPACK-mediated lipidation (**Fig 6C**). SDS-PAGE of XP-L115R released by detergent solubilization migrated slowly and remained near the top of the gel, suggesting XPACK induced high-order oligomerization or aggregation (**Fig 6C**). In spite of the apparent difference between DNAJC5 XP-L115R and WT DNAJC5, α-syn secretion was stimulated by the expression of both species (**Fig 6D**). Treatment with the palmitoylation inhibitor 2-BA resulted in the formation of XP-L115R that migrated to a position similar to that of monomeric DNAJC5 (**Fig 6E**). Correspondingly, secretion of α-syn was no longer stimulated by the palmitoylation deficient DNAJC5 XP-L115R monomer (**Fig 6D**).

In order to expand on the fractionation results, we employed confocal microscopy to examine the subcellular localization of DNAJC5 XP-L115R fused with a C-terminal HaloTag. In contrast to the diffuse distribution of the DNAJC5 L115R mutant, which was excluded from the interior of enlarged endosomes (**Fig 4 supplement 2A**), punctate DNAJC5 XP-L115R was widely associated with enlarged endosomes (**Fig 6 supplement 2A**). Internalization events were found in several enlarged endosomes (**Fig 6 supplement 2A, magnified insets**). mNG-α-syn was also incorporated into endosomal compartments in cells co-expressing DNAJC5 XP-L115R (**Fig 6 supplement 2B**). The level of α-syn containing endosomes in cells expressing DNAJC5 XP-L115R was ∼2-fold higher than in cells expressing the DNAJC5 L115R mutant (**Fig 6 supplement 2C**). Our imaging data corroborate the biochemical similarity between WT DNAJC5 and DNAJC5 XP-L115R.

### Secretion of endogenous α-syn from neurons is mediated by DNAJC5

To evaluate the function of DNAJC5 in α-syn secretion at physiological levels of expression in a neuronal cell line, we employed SH-SY5Y, a neuroblastoma line that differentiates in the presence of retinoic acid (RA) into nerve cells that express dopamine neuron (DA) markers including tyrosine hydroxylase (TH) (Lopes et al., 2010). We observed elevated levels of expression of α-syn and dopamine transporter (DAT) in SH-SY5Y cells after six days of RA-induced differentiation (**Fig 7 supplement 1A**). Fractionation of SH-SY5Y cell lysates resolved DNAJC5 into the low electrophoretic mobility P form associated with sedimentable membranes (M) and the non-sedimentable cytosolic NP form (**Fig 7A**). Hydroxylamine treatment of the membrane associated form converted DNAJC5 to the electrophoretic mobility position of the NP form, as before (**Fig 7 supplement 1B; Fig. 1 supplement 1B**).

**Figure 7.**
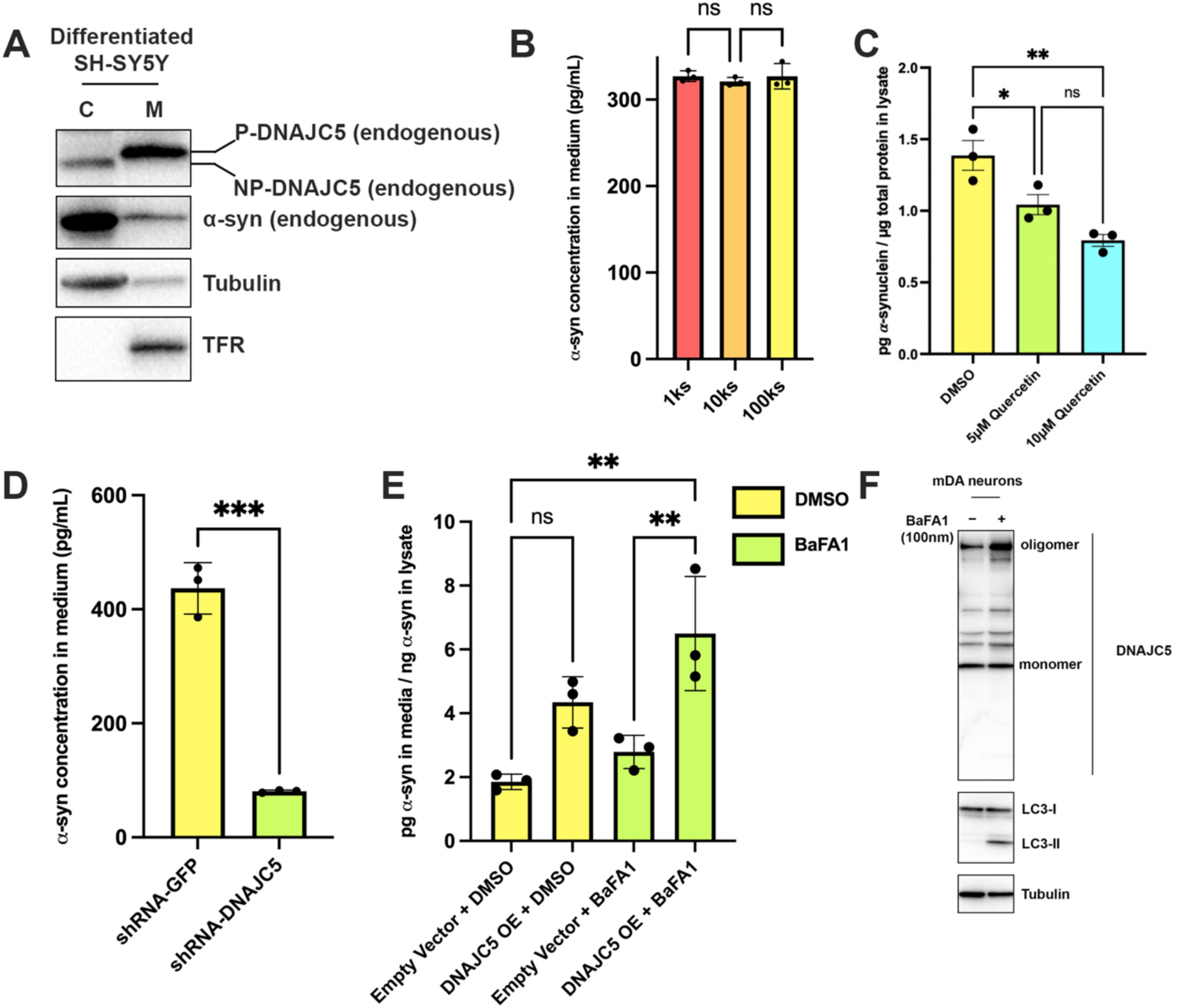
Recapitulation of endogenous DNAJC5-mediated α-syn secretion in various neuronal cell cultures. (A) Membrane and cytosol fractionation of differentiated SH-SY5Y neuroblastoma cells. The fractionation was performed as depicted in Figure 1B. C, cytosol. M, membrane. The distribution of endogenous DNAJC5 and α-syn was evaluated by immunoblot. Transferrin receptor (TFR) was used as a membrane marker. Tubulin was used as a cytosol marker. (B) Quantification of α-syn level in the supernatant of centrifuged media with ELISA. Conditioned media were collected and sequentially centrifuged at 1,000 (1k) g, 10,000 (10k) g, and 100,000 (100k) g. The supernatant from each centrifugation step (1ks, 10ks and 100ks) was collected and measured by LEGEND MAX^TM^ Human α-synuclein (Colorimetric) ELISA Kit. One-way ANOVA showed no significant (ns) difference of α-syn level between fractions. (C) Quercetin inhibited endogenous α-syn secretion in hiPSC-derived midbrain dopamine neurons. hiPSC-dopamine neurons carrying the *GBA-N370S* mutation were treated with quercetin (5 μM or 10 μM) at Day 35. Culture media samples were harvested after 3d treatment at Day 38 and α-syn levels in the media were analyzed by electro-chemiluminescent immunoassay. Data points represent individual cell lines derived from different donors and are normalised to total protein in the corresponding cell lysates. One-way ANOVA followed by Tukey’s post-hoc test shows a significant reduction in α-syn secretion with increasing quercetin concentration (*p<0.05, **p<0.01). (D) Depletion of endogenous DNAJC5 in SH-SY5Y cells decreased basal α-syn secretion. After 3d of culture, the media from differentiated SH-SY5Y cells transduced with shRNA targeting GFP (shRNA-GFP) or shRNA targeting DNAJC5 (shRNA-DNAJC5) were collected and the extracellular α-syn was quantified with ELISA. ***p value<0.0002, two-tailed t test. (E) Expression of exogenous human DNAJC5 in mouse mDA stimulated basal α-syn secretion. WT mDA and mDA expressing hDNAJC5 were treated with DMSO or 100 nM BaFA1. Quantification of α-syn in conditioned media was performed with Mouse α-synuclein ELISA Kit (Abcam). α-syn secretion was normalized by dividing the α-syn in media (pg/mL) by the α-syn in cell lysates (ng/mL). **p value<0.01, one-way ANOVA. (F) BaFA1 increased DNAJC5 oligomerization in mouse mDA neurons.

We employed a sensitive α-syn enzyme-linked immunosorbent assay (ELISA) and detected about 300 pg/ml α-syn secreted into the supernatant of RA-differentiated SH-SY5Y cells (Forland et al., 2018) (**Fig 7B**). Differential centrifugation of the medium fraction demonstrated that the bulk of the secreted α-syn remained soluble (**Fig 7B**). A 100k pellet fraction was probed by protease protection for the localization of Flot-2, an exosome marker and DNAJC5. Both were resistant to degradation by proteinase K in the absence but not in the presence of TX-100, suggesting that both were protected within the lumen of EVs (**Fig 7 supplement 1C**). In contrast, residual full-length (FL) α-syn in the pellet fraction was cleaved by proteinase K without or with detergent (**Fig 7 supplement 1C**). As an additional test, sucrose gradient flotation of the high-speed pellet fraction as used in **Fig 2 supplement 2A,** revealed that α-syn secreted by differentiated SH-SY5Y cells was not buoyant whereas DNAJC5 appeared associated with membranes fractionating at the position of EVs (**Fig 7 supplement 1D**). Thus, as with HEK293T cells, α-syn secreted by differentiated SH-SY5Y cells appears not to be enclosed within EVs.

Midbrain dopamine (mDA) neurons (**Fig 7 supplement 2A**) differentiated from human induced pluripotent stem cells (hiPSCs) from Parkinson’s patients with the *GBA-N370S* mutation, a genetic lesion which causes ER stress and dysfunctional lysosomes, release about twice the level of α-syn compared to control WT neurons(Fernandes et al., 2016; Lang et al., 2019) Treatment of *GBA-N370S* hiPSC-derived dopamine neurons with the DNAJC5 inhibitor quercetin led to a significant dose-dependent reduction in the secretion of endogenous α-syn (**Fig 7C**). Immunnoblotting of hiPSC-derived dopamine neuron lysate revealed that the endogenous DNAJC5 is natively palmitoylated which can be partially reduced by treatment with 2-BA (10 µM) to induce the formation of the lower, non-palmitoylated band (**Fig 7 supplement 2B**). However, this partial de-palmitoylation of DNAJC5 was insufficient to inhibit α-syn release by iPSC-derived *GBA-N370S* dopamine neurons at the concentration of 2-BA used (**Fig 7 supplement 2C**).

To examine the role of DNAJC5 in differentiated SH-SY5Y cells, we silenced the expression of the chromosomal locus by small hairpin RNAs (shRNAs) transduced by lentivirus. The efficiency of shRNA targeting DNAJC5 was confirmed by knockdown (KD) of endogenous DNAJC5 in HEK293T cells (**Fig 7 supplement 3A**). Similarly, DNAJC5 was successfully depleted in differentiated SH-SY5Y cells (**Fig 7 supplement 3B**). As a result, secretion of α-syn was reduced 5-fold compared with a control transduced with shRNA targeting GFP (**Fig 7D**).

In HEK293T cells, overexpression of DNAJC5 increased α-syn secretion (**Fig 1E**). We were unable to observe enhanced secretion of α-syn in SH-SY5Y overexpressing DNAJC5, possibly because of a high level of expression of endogenous DNAJC5 in the differentiated cells (**data not shown**). To test the effect of DNAJC5 on the basal level of α-syn secretion in neurons, we stably transduced mouse embryonic stem cells (mESCs) with lentivirus containing human DNAJC5 WT or L115R and differentiated them into mDA neurons (**Fig 7 supplement 4A**). After differentiation, DNAJC5 WT was expressed in mDA, but we were unable to detect the expression of DNAJC5 L115R (**Fig 7 supplement 4B**). Analysis of conditioned media by ELISA revealed a two-fold elevated α-syn secretion in DNAJC5 WT overexpressing mDA compared to control with an empty vector in the presence of BaFA1 (**Fig 7E**). With both BaFA1 treatment and DNAJC5 overexpression, α-syn secretion was increased three fold (**Fig 7E**). We examined the cell lysate of mDA by immunoblot. The effect of BaFA1 inhibition was indicated by the appearance of a lipidated form of LC3 (LC3-II) (**Fig 7F**). BaFA1 treatment also induced more DNAJC5 oligomer formation (**Fig 7F**). We conclude that DNAJC5 stimulates α-syn secretion in differentiated DA neurons as it does in HEK293T cells.

### The J domain and C-terminal tail (C tail) of DNAJC5 are dispensable for α-syn secretion

The secretion deficiency caused by the L115R mutation highlights the importance of the CS domain of DNAJC5 in regulating α-syn secretion. The structure of the J domain of DNAJC5 has been solved by nuclear magnetic resonance (NMR) (Patel et al., 2016). Recent progress in deep learning algorithms, exemplified by AlphaFold, enables atomic accuracy in protein structure prediction (Jumper et al., 2021). We searched the public AlphaFold database to examine the predicted structure of full-length DNAJC5. In the predicted structure, the J domain showed a conserved overall globular J protein fold within the N-terminus, linked by the helical CS domain to the flexible C-terminal tail. Only a short helix was predicted to reside within the C-tail (**Fig 8A**). We refined the boundary of each domain in DNAJC5 based on the predicted structure.

**Figure 8.**
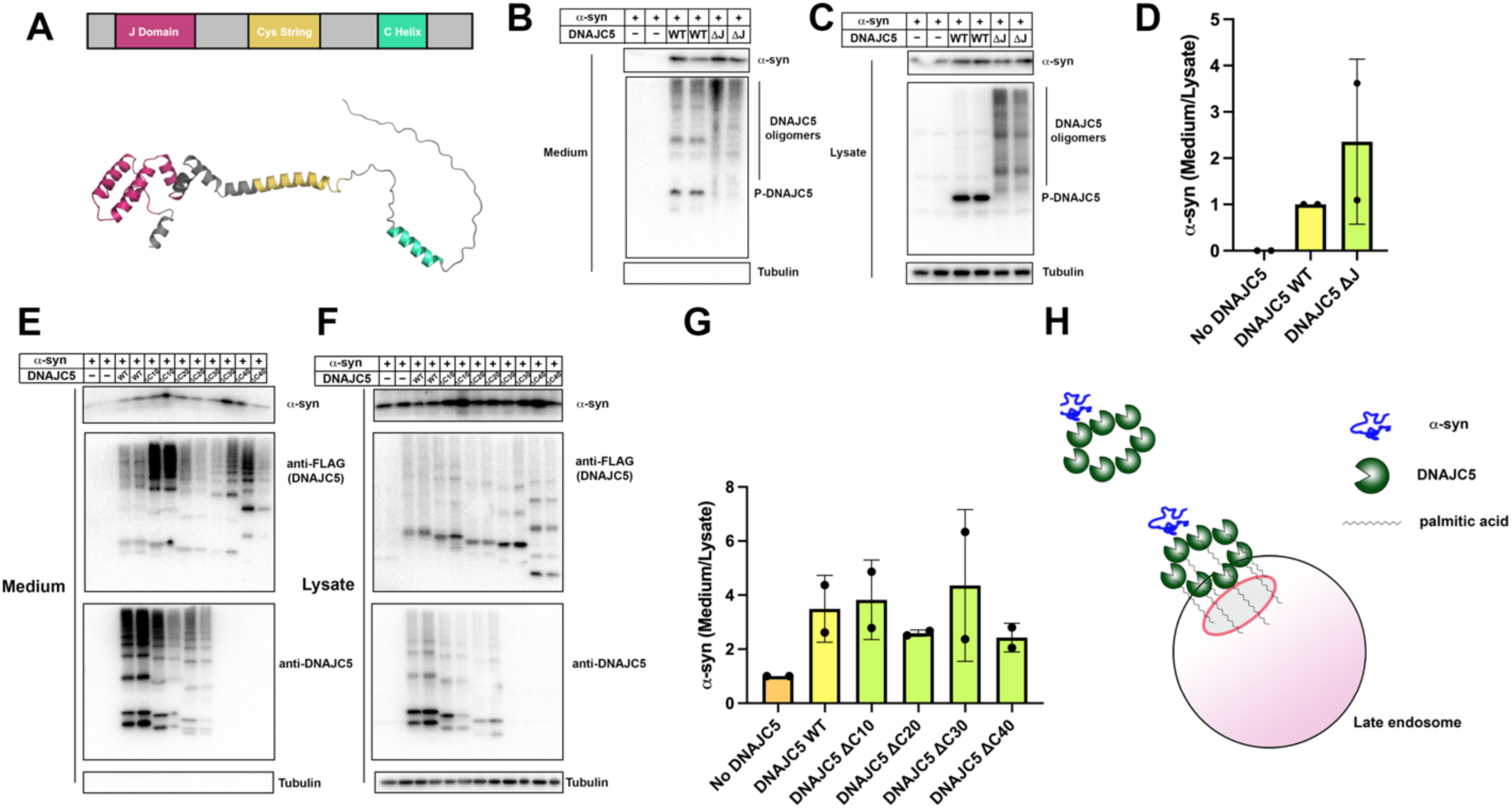
Domain mapping of secretion-competent DNAJC5. (A) Predicted structure of DNAJC5 by AlphaFold. Color scheme: J domain (magenta), Cys string domain (yellow) and C-terminal helix (green). (B) DNAJC5 (ΔJ) was competent to induce α-syn secretion into the medium. HEK293T cells were transfected with indicated plasmids. Media were collected after 36 h and evaluated with immunoblot. (C) DNAJC5 (ΔJ) formed oligomers in HEK293T cells. (D) Quantification of normalized α-syn secretion in HEK293T cells transfected with WT DNAJC5 or DNAJC5 (ΔJ). Quantification was based on immunoblot in (B) and (C). The α-syn secretion was calculated as the amount of α-syn in media divided by the amount in lysate. α-syn secretion in cells transfected with WT DNAJC5 was normalized as 1. (E) C-terminal truncated DNAJC5 constructs were competent to induce α-syn secretion in the medium. HEK293T cells were transfected with C-terminal truncated DNAJC5 and α-syn. DNAJC5 antibodies cannot recognize DNAJC5 (ΔC30) and DNAJC5 (ΔC40) because of a missing epitope in the C-terminus. Instead, DNAJC5 (ΔC30) and DNAJC5 (ΔC40) were detected by C-terminal FLAG tags. All the C-terminal truncated DNAJC5 constructs showed smear-like oligomers. (F) Expression of C-terminal truncated DNAJC5 constructs in HEK293T cells. Immunoblot of anti-FLAG antibody and anti-DNAJC5 antibody cross-validated the existence of oligomers. (G) Quantification of normalized α-syn secretion in HEK293T cells transfected with WT DNAJC5 or different C-terminal truncated DNAJC5 constructs (ΔC10, ΔC20, ΔC30 and ΔC40). Quantification was based on immunoblot in (E) and (F). The α-syn secretion was calculated as the amount of α-syn in media divided by the amount in lysate. α-syn secretion in cells without DNAJC5 transfection was normalized as 1. (H) A model for palmitoylated DNAJC5 oligomer-mediated α-syn secretion.

Using this information, we showed that oligomerization of DNAJC5 increased when the J domain was deleted (**Fig 6 supplement 1A and Fig 8C**). As recently reported by Lee et al. (2022), deletion of the J domain increased the level of α-syn secretion induced by DNAJC5 (**Fig 8B&8D**). Next, we examined the function of the C-tail by truncating about ten amino acids at a time, resulting in a series of C-terminal truncated DNAJC5 constructs, i.e., DNAJC5 ΔC10, ΔC20, ΔC30, ΔC40. All four DNAJC5 C-terminal truncations were expressed and formed oligomers in cells (**Fig 8F**). C-terminal truncated oligomers were co-secreted with α-syn into the medium (**Fig 8E&8G**). This result demonstrates that neither the J domain nor the C-tail is required for DNAJC5 to induce α-syn secretion.

## Discussion

Transmission of protein aggregates and subsequent self-amplification is emerging as a common theme across various neurodegenerative diseases. DNAJC5 has been shown to control the release of neurodegenerative disease proteins but the mechanism of action of this protein in unconventional secretion remains elusive. In this study, we reconstituted DNAJC5-regulated α-syn secretion in cultured HEK293T cells, in RA neuronally differentiated human cells and in hiPSC-derived midbrain DA neurons. By combining this assay with medium and cellular fractionation, we demonstrated that membrane-anchoring of DNAJC5 through palmitoylation is crucial for its secretion and the secretion of α-syn as a soluble monomer. In addition, we observed the topological locations of DNAJC5 and α-syn within enlarged endosomes, presumably at an intermediate stage prior to secretion. Furthermore, DNAJC5 was found to form oligomers and the importance of the oligomerzation was highlighted by the use of a lipidated XPACK fusion peptide. Our findings on the role of DNAJC5 extend to differentiated DA neurons of human and mouse origin. Finally, we provide evidence that both palmitoylation and oligomerization are solely dependent on the CS domain, which is required for α-syn secretion. Based on our biochemical assays and imaging observations, we propose that palmitoylated DNAJC5 oligomers function at a step involving membrane translocation of cytosolic α-syn, enabling it to become competent for secretion (**Fig 8F**).

The in vivo toxicity of α-syn aggregates remains elusive (Lashuel et al., 2013) but its propagation accompanies the progression of PD (Braak et al., 2006; Braak et al., 2003). Recently, (Calo et al., 2021) found that DNAJC5 expression decreases in α-syn transgenic mice. Overexpression of DNAJC5 in vivo is reported to rescue α-syn aggregation-dependent pathology and increase the accumulation of monomeric α-syn (Calo et al., 2021). As we find and others have reported, iPSC-derived neurons also secrete α-syn in a largely soluble form (Fernandes et al., 2016). Using the criteria of differential sedimentation and gel filtration chromatography, we conclude that α-syn is secreted in cultures cells as a soluble monomeric species not enclosed within extracellular vesicles, regardless of mutations modeled on PD (**Fig 2 supplement 1A**) or as expressed in tandem arrays or in gene fusions to tightly folded proteins (**Fig 5 supplement 1**). Consistent with our results, other MAPS substrates are also reported to be secreted in a soluble form (Lee et al., 2016). Although α-syn oligomers have also been found in EVs (Danzer et al., 2012; Emmanouilidou et al., 2010; Guo et al., 2020) we see no evidence for this in our culture medium fractionation and immunoblot experiments (**Fig 2B**). With a more sensitive and quantitative Nluc assay, 85% of secreted α-syn was found to be soluble (**Fig 2 supplement 1B**). Similarly, the basal level of α-syn secreted by differentiated neuroblastoma cells is mainly soluble and not detected within EVs (**Fig 7 supplement 1A**).

The release of soluble α-syn may be an early event in pathogenesis of PD, prior to the deposition of aggregates. Inhibition of the lysosomal ATPase with bafilomycin A (BaFA1) is known to induce lysosome fusion to the cell surface and secretion of lysosomal content including both soluble and aggregate forms of α-syn (Xie et al., 2022) **(Fig 1 supplement 3H, Fig 7E**). Such secretion may be part of a cell protective mechanism but it may also promote the interneuronal spread of monomer and oligomer.

Numerous neuronal proteins are palmitoylated, including synaptic scaffolding proteins, signaling proteins and synaptic vesicle proteins (Linder and Deschenes, 2007). Protein palmitoylation has been implicated in the pathogenesis of neurodegenerative diseases (Cho and Park, 2016). In PD particularly, a recent study reported that upregulation of cellular palmitoylation decreased α-syn cytoplasmic inclusions (Ho et al., 2021). The neuropathology and behavior deficiency of Huntington disease (HD) mice can be reversed by boosting brain palmitoylation (Virlogeux et al., 2021). In the case of DNAJC5, L115R and L116Δ, the two adjacent mutations causing decreased palmitoylation of DNAJC5 monomers, lead to a familial form of NCL (Benitez et al., 2011; Diez-Ardanuy et al., 2017). Our results suggested that the secretion of neurodegenerative disease proteins is also dependent on palmitoylation, possibly alleviating the cellular burden of protein aggregate accumulation. Notably, the general inhibition of cellular palmitoylation by 2-BA led to a complete block of α-syn secretion, whereas the specific palmitoylation deficient DNAJC5 mutant L115R only partially decreased α-syn secretion (**Fig 3**). The difference implies the existence of palmitoylation-dependent factors other than DNAJC5.

Although DNAJC5 co-expressed with α-syn and palmitoylation are required for secretion, EV-associated P-DNAJC5 clearly separated from soluble α-syn in the culture medium (**Fig 2B and Fig 2 supplement 2B**). At which step do the two separate? In our live cell imaging experiments, the internalized DNAJC5 inside enlarged endosomes had both punctate and diffuse distributions (**Fig 4A**). This may represent the soluble NP-DNAJC5 and membrane-attached P-DNAJC5, respectively. In the time-lapse imaging of internalized α-syn induced by DNAJC5, both DNAJC5 and α-syn moved dynamically inside the compartment, without significant co-localization (**Fig4 supplementary video**). This observation suggests that the separation of DNAJC5 and α-syn may occur prior to their secretion when the late endosome and plasma membrane fuse.

Zhang et al. (2020) have reported a novel membrane channel, TMED10, for the unconventional secretion of IL-1β. These authors speculate an activation-on-demand oligomerization of TMED10 membrane subunits to form a conducting channel for substrate translocation, a process they refer to as THU (Zhang et al., 2020). α-syn was reported to not depend on TMED10 for unconventional secretion (Zhang et al., 2020). Similarly, in chaperone-mediated autophagy (CMA), cytosolic substrates are proposed to be translocated into lysosomes through a channel formed by the oligomerization of a single-transmembrane protein, LAMP2A (Bandyopadhyay et al., 2008). The secretion of α-syn has been shown to be independent of CMA (Lee et al., 2018). Without a transmembrane domain, membrane-tethered P-DNAJC5 oligomer is unlikely to be a channel for translocation. A recent report identified CD98hc, an amino acid transporter subunit, to be a DNAJC5 interactor that is required for α-syn secretion (Lee et al., 2022). It remains to be determined whether CD98hc or other as yet uncharacterized membrane proteins are directly involved in α-syn membrane translocation.

In SEC61-mediated co-translational translocation, substrates enter the SEC61 translocon in an unfolded state (Rapoport et al., 2017). In THU and CMA, substrate unfolding is also required for translocation across the membrane (Kaushik and Cuervo, 2018; Zhang et al., 2020). In striking contrast, unfolding and size is not a limiting factor for α-syn secretion (**Fig 5**). As a precedent of translocation without unfolding, studies have shown the import of folded proteins into the matrix of peroxisomes and obviously through the nuclear pore (Kim and Hettema, 2015; Lin and Hoelz, 2019). DNAJC5 forms a series of extremely stable oligomers, which may provide versatile adaptors to accommodate diverse misfolded or folded substrates with different dimensions. The structure of DNAJC5 oligomers may shed light on the principle of this folding-independent translocation pathway.

## Materials and methods

### Key Resources table

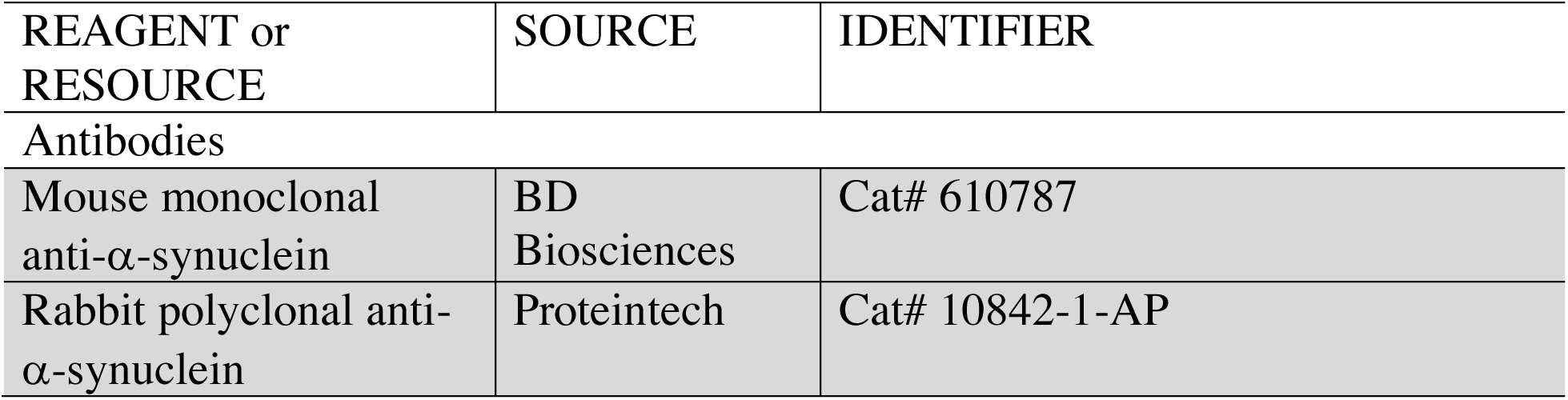
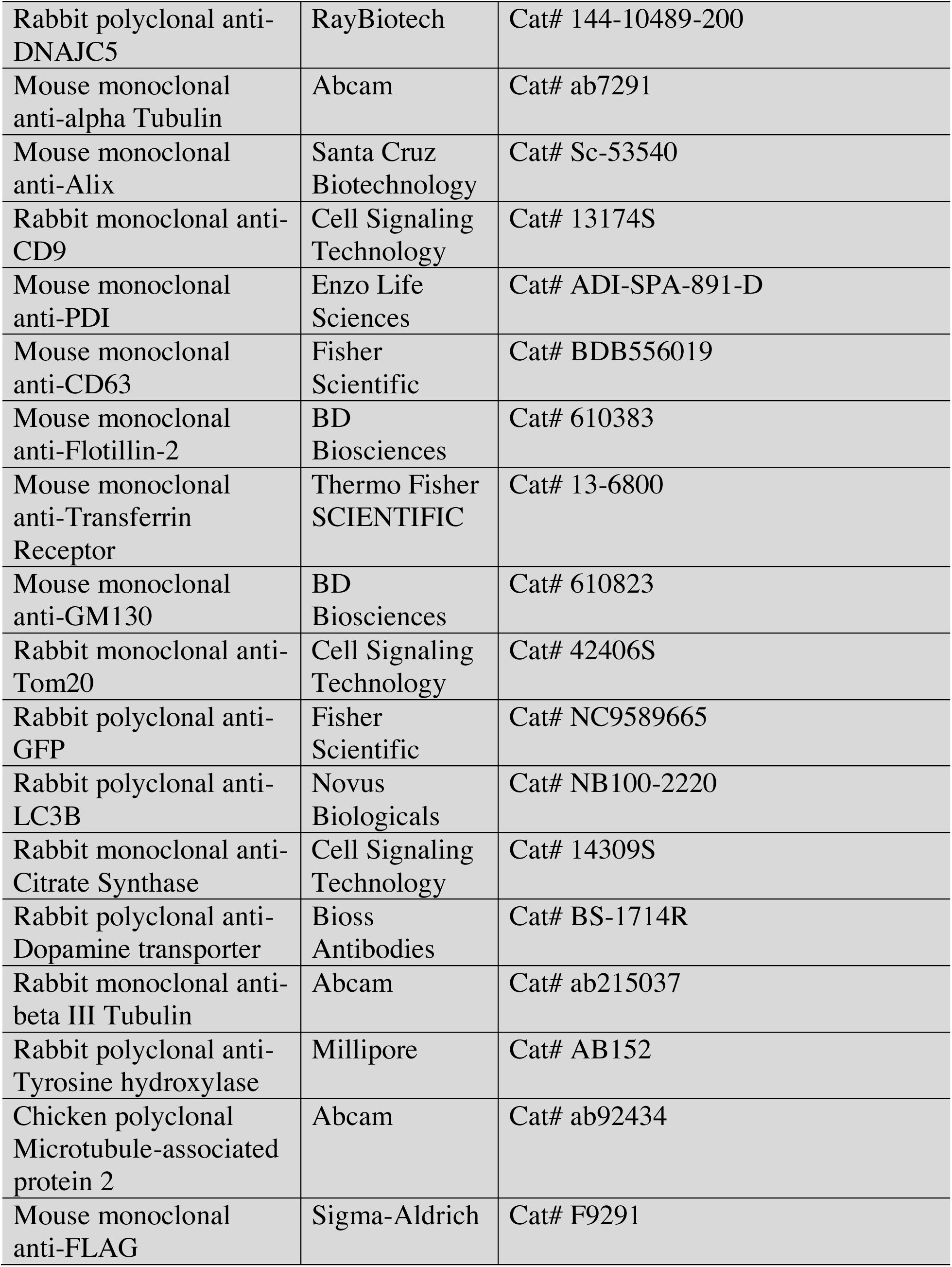
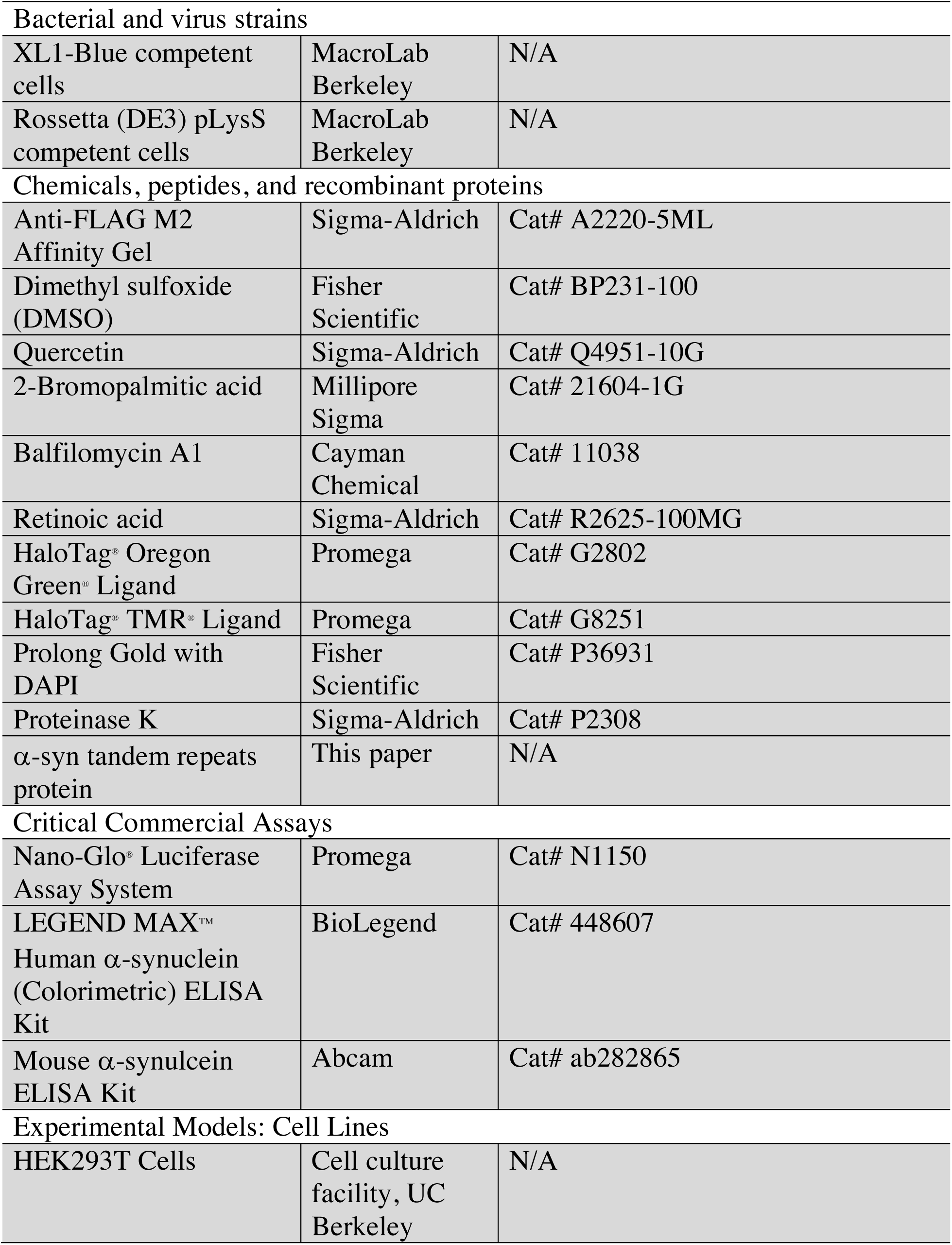
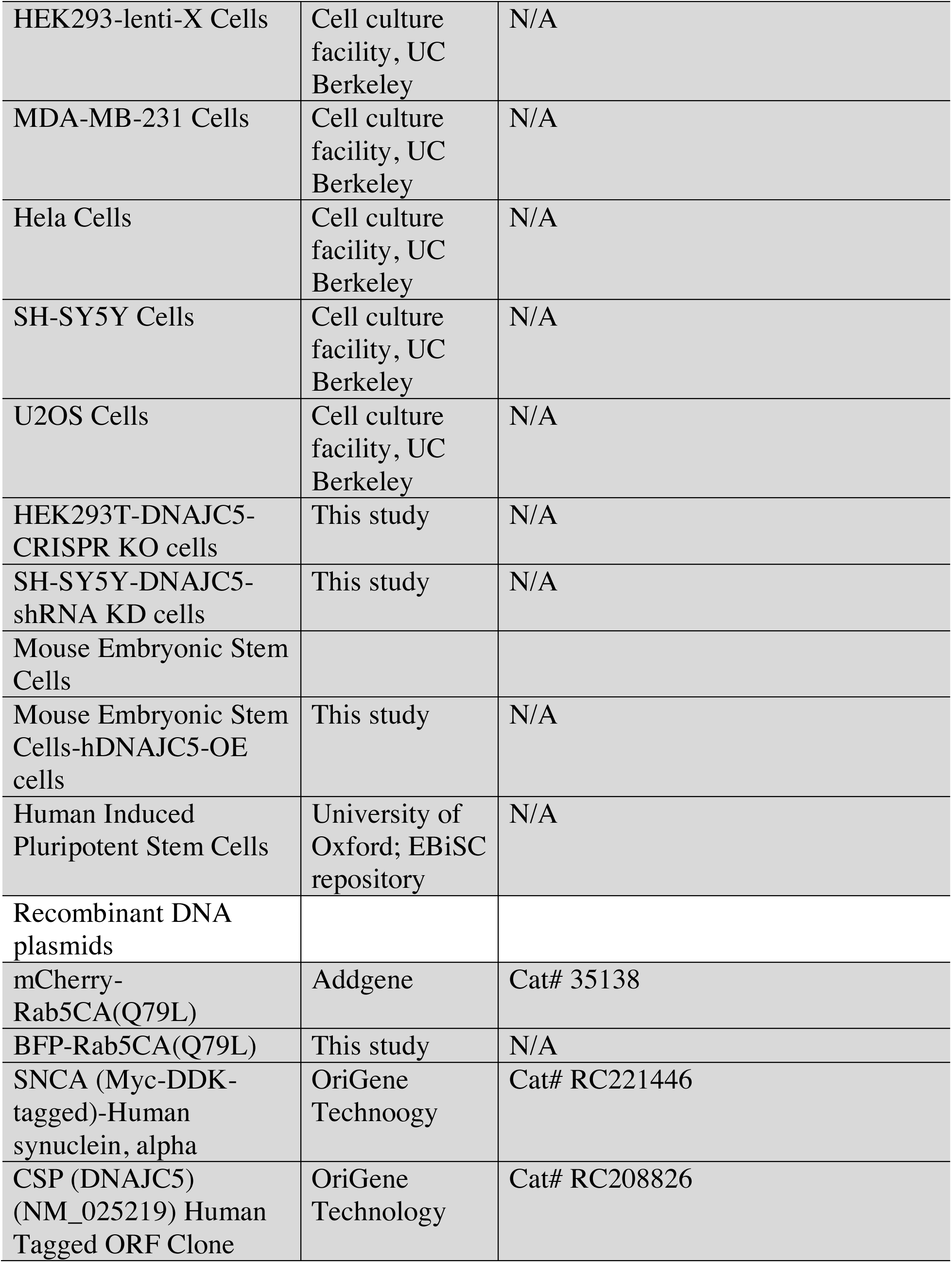
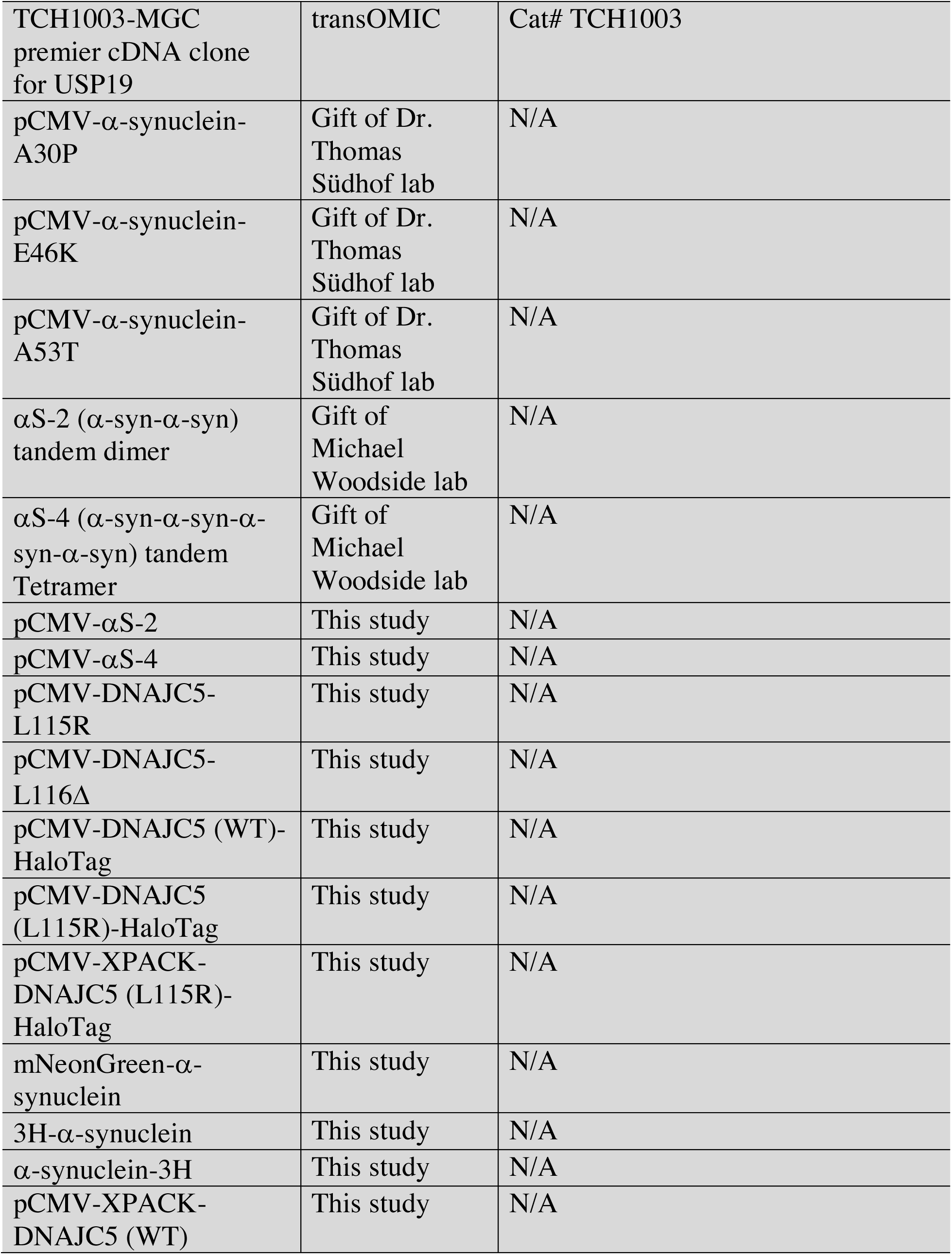
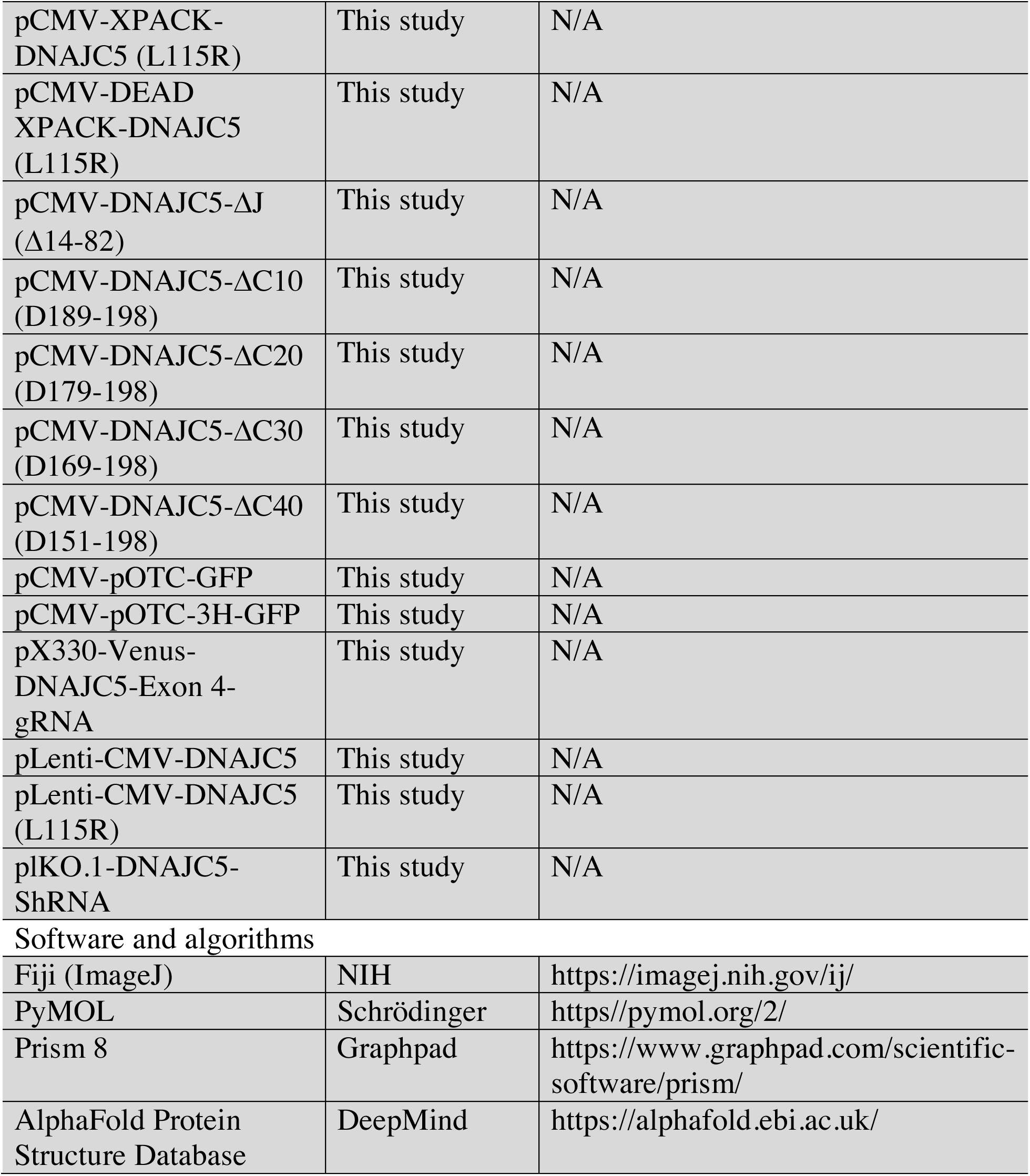

### Cell culture and transfection

Cells were grown at 37°C in 5% CO2 and maintained in Dulbecco’s modified Eagle’s medium (DMEM) supplemented with 10% fetal bovine serum (FBS). For secretion assays, the FBS concentration was reduced to 1% for up to 36 h during which time the growth rate of cells slowed but cells remained viable. For EV preparation and medium fractionation, we grew cells in DMEM supplemented with exosome-depleted FBS. Exosome-depleted FBS was prepared by overnight centrifugation of 30% diluted FBS in DMEM at 100,000xg. Transfection of plasmids into cells was performed using Lipofectamine 2000 (Thermo Fisher Scientific, Waltham, MA) according to the manufacturer’s protocols.

### Reconstitution of α-syn secretion in HEK293T cells

HEK293T cells (cultured in 6-well plates) were cultured to 60% confluence and co-transfected with plasmids encoding different constructs of α-syn and DNAJC5. pCMV-GFP was used as a transfection control in all secretion experiments. At 4 h after transfection, we replaced cell culture medium with DMEM supplemented with 1% FBS containing indicated drugs for treatment. At indicated time points, we collected media fractions which were centrifuged at 1,000xg for 10 min to remove floating cells and cell debris. The media were mixed with methanol/chloroform to precipitate proteins which were collected by centrifugation (10,000 g x 10 min) and resuspended in SDS-PAGE sample loading buffer to achieve concentration (20-fold). Cells were lysed in lysis buffer (10 mM Tris, pH 7.4, 100 mM NaCl, 1% Triton X-100). Both the concentrated media and cell lysate fractions were analyzed by immunoblot.

To exclude the release of cytoplasmic proteins from cell death, we monitored the viability of cells after transfection with a Countess^TM^ II Automated Cell Counter (Thermo Scientific) using trypan blue staining.

For a nanoluciferase-based assay, media fractions were collected and centrifuged at 1,000xg for 10 min. The supernatant fractions were harvested and further diluted with PBS buffer (1000-fold). The nanoluciferase activity was assayed using a Nano-Glo^®^ Luciferase Assay System (Promega, Madison, WI) according to the manufacturer’s protocol.

### Membrane and Cytosol fractionation

Cells (one 10cm dish) were cultured to 70% confluence and transfected with different constructs of DNAJC5. One day after transfection, we harvested the transfected cells by scraping in 1 ml B88 (20 mM HEPES-KOH, pH 7.2, 250 mM sorbitol, 150 mM potassium acetate and 5 mM magnesium acetate) plus a cocktail of protease inhibitors (Sigma-Aldrich, St. Louis, MO). Cells were homogenized by 10 passages through a 22G needle. Homogenates were centrifuged at 500xg for 10 min and the resulting post-nuclear supernatant (PNS) fractions were centrifuged at 100,000xg for 1.5 h. High-speed supernatant fractions were then subjected to a repeat centrifugation to achieve a clarified cytosol fraction. The pellet fraction was washed and resuspended in the same volume of B88. Resuspended material was also centrifuged again to collect a washed membrane fraction. Membranes were lysed in lysis buffer.

For membrane fractionation, the PNS was subjected to differential centrifugation at 3,000xg (10 min), 25,000xg (20 min) and 100,000xg (30min). Membrane fractions were normalized to phosphatidylcholine content and analyzed by immunoblot (Ge et al., 2013).

For proteinase K protection assays, the 25,000xg membrane fraction was aliquoted into three tubes: one without proteinase K, one with proteinase K (10 μg/ml), and one with proteinase K plus TritonX-100 (0.5%). The incubation was conducted on ice for 20 min and stopped by sequential addition of PMSF (1mM) and sample buffer and samples were then heated on metal block at 95°C for 5 min and analyzed by SDS-PAGE and immunoblot.

### In vitro depalmitoylation assay

Cells (HEK293T, MDA-MB-231 or Hela) were transfected with DNAJC5. Cellular membranes were prepared as described above. For chemical deplamitoylation, the membranes were resuspended and incubated with 0.5 M hydroxylamine (pH 7.2) or 0.5 M Tris (pH 7.2, control) at room temperature overnight in the presence of a cocktail of protease inhibitors (Sigma-Aldrich, St. Louis, MO). The mobility of DNAJC5 was examined by SDS-PAGE followed by immunoblot.

### CRISPR/Cas9 genome editing

gRNA targeting exon 4 of DNAJC5 (CACCGGAGGCCGCAGAAGACAAACA) was inserted into a pX330-based plasmid expressing Venus fluorescent protein (Shurtleff et al., 2016). HEK293T cells were transfected with pX330-pX330-Venus-DNAJC5-Exon 4-gRNA by Lipofectamine 2000 (Thermo Fisher Scientific, Waltham, MA). After 48 h, we diluted the cells and single colonies were isolated, expanded and determined for DNAJC5 KO by immunoblot.

### Medium fractionation and extracellular vesicle preparation

Conditioned medium was harvested and centrifuged first at 1,500xg for 20 min followed by 10,000xg for 30 min and 100,000xg for 1.5 h. The supernatant fractions at each step were collected and treated with methanol/chloroform to precipitate proteins which were then collected by centrifugation. Pellet fractions were resuspended in sample buffer to achieve a 20-fold concentration. The sedimented fractions at each step were also collected and resuspended in sample buffer. All the fractions were analyzed by immunoblot.

EVs were isolated by buoyant density flotation on a sucrose step gradient. The pellet fraction from a 100,000xg centrifugation was resuspended in PBS and mixed with 60% sucrose buffer (10mM Tris-HCl pH 7.4, 100 mM NaCl) to achieve a final sucrose concentration >50% as measured with a refractometer. Aliquots of 40% (5 ml) and 10% (2 ml) sucrose buffer were sequentially overlaid above the sample. The tubes were then centrifuged at 150,000xg for 16 h in an SW41 Ti swinging-bucket rotor (Beckman Coulter). After centrifugation, 0.5 ml fractions were collected from top to bottom and samples analyzed by SDS-PAGE and immunoblot.

### Co-immunoprecipitation

Media fractions were collected and centrifuged at 1,000xg for 10min. The supernatant fractions were collected and concentrated (20-fold) using a 10 kDa Amicon filter (Millipore, Billerica, MA). Concentrated media fractions (1 ml) were incubated with 20 μl of anti-FLAG M2 affinity gel (Sigma-Aldrich, St. Louis, MO) for 1h at 4°C. After washing 5x with lysis buffer, SDS-PAGE sample loading buffer was added to the beads and samples were processed for SDS-PAGE and immunoblot.

### Protein purification

The purification of different α-syn tandem-oligomer constructs was performed as previously described (Dong et al., 2018). Briefly, an osmotic shock protocol was adapted to enrich proteins released from the periplasm of transfected *E. coli*. The supernatant fraction containing released proteins was subjected to ammonium sulfate (AS) precipitation, with 50%, 45% and 40% saturated concentration of AS for monomer, dimer and tetramer, respectively. After overnight precipitation, the precipitated proteins were collected by centrifugation at 100,000xg for 30 min. The pellet fractions were dissolved in Buffer A (20mM Tris-HCl pH 8.0) and clarified by repeated centrifugation at 100,000xg for 30 min.

Clarified supernatants were applied to an equilibrated HiPrep Q Fast Flow 16/10 column (GE Healthcare, Chicago, IL). Eluted proteins were collected, concentrated by 10 kDa Amicon filter (Millipore, Billerica, MA) and further purified by gel filtration (Superdex-200, GE Healthcare) with PBS used as gel filtration buffer. Purified proteins were assessed by SDS-PAGE followed by coomassie-blue staining.

### Immunofluorescence and live-cell Imaging

For immunofluorescence, U2OS cells were washed once with PBS and immediately fixed by 4% EM-grade paraformaldehyde (Electron Microscopy Science, Hatfield, PA) for 10 min at room temperature. Cells were washed three times with PBS and blocked and permeabilized for 30 min in permeabilization buffer (5% FBS and 0.1% saponin in PBS). hiPSC dopamine neurons were fixed with 4% paraformaldehyde in PBS and 0.1% Triton-X was used for permablization (10 mins) followed by blocking in 10% normal donkey serum for 1 h. Cells were then incubated with 1:100 dilution of primary antibodies overnight at 4°C. After three washes with PBS, cells were incubated with 1:500 dilution of fluorophore-conjugated secondary antibodies for 30 min at room temperature. Prolong Gold with DAPI (Thermo Fisher) was used as mounting solution. Images were acquired with a Zeiss LSM900 confocal microscope and analyzed with Fiji/ImageJ software (https://imagej.nih.gov/ij/).

For live-cell imaging, cells were cultured in 35mm glass bottom dishes (MatTek). The addition of HaloTag fluorescent ligands were added according to the manufacturer’s protocol (Promega). After incubation, the medium was replaced with Opti-MEM supplemented with 10% FBS. Imaging was performed using a Zeiss LSM900 confocal microscope in a temperature-controlled (37°C and 5% CO2) environment.

### Mitochondria purification

HEK293T cells were trypsinized and collected by centrifugation. Cells were washed twice with NKM buffer (1 mM Tris HCl, pH7.3, 0.13 M NaCl, 5mM KCl, 7.5 mM MgCl_2_), and resuspended in 6 packed cell volumes of homogenization buffer (10 mM Tris pH 7.4, 10 mM KCl, 0.15 mM MgCl_2_). Cells were homogenized by 10 passages through a 22G needle. Cell homogenates were mixed gently with the same volume of 2.3M sucrose solution and centrifuged at 1,200xg for 5 min to remove unbroken cells and large cell debris. The recovered supernatant fractions were centrifuged at 7,000xg for 10min. Mitochondria enriched in the pellet fraction were resuspended in 3 packed cell volumes of Mitochondria Suspension Buffer (10 mM Tris, pH 7.3, 0.15 mM MgCl2, 0.25mM sucrose).

### Differentiation of SH-SY5Y cells

SH-SY5Y neuroblastoma cells were maintained in Dulbecco’s modified Eagle’s medium (DMEM) supplemented with 1x non-essential amino acid (NEAA), 1x sodium pyruvate and 10% fetal bovine serum (FBS). Differentiation was induced by lowering the FBS in culture medium to 1% plus 10 μM retinoic acid (RA). Cell medium was replaced each 3 d to replenish RA. Cell morphology was monitored by microscopy and experiments on SH-SY5Y cells were performed from D6 of differentiation.

### shRNA knockdown

plKO.1-Hygro plasmids containing shRNA targeting DNAJC5 (ccggGCAACCTCAGATGACATTAAACTCGAGTTTAATGTCATCTGAGGTT GCTTTTTG) together with pMD2.G and PsPAX2 were transfected into HEK293T cells to produce lentiviral particles for 72 h. Lentivirus particles were concentrated with Lenti-X^TM^ Concentrator (Takara Bio). SH-SY5Y was transduced by lentivirus before differentiation. Three days post transduction, cells were selected with 250 μg/ml hygromycin for 10 d. The selected cells were differentiated, and the knockdown was verified with immunoblot.

### Culture and differentiation of mouse embryonic stem cells

Mouse ESCs (R1) were maintained and differentiated into dopaminergic neurons following a modified protocol from (Ni et al., 2013). Briefly, R1 cells were maintained in a feeder-independent system, plated in 0.1% gelatin (StemCell technologies) and cultured in KSR medium consisting of KnockOut DMEM, 20% KnockOut serum replacement, 2mM L-glutamine, 0.1mM nonessential amino acids, 0.1 mM β-mercaptoethanol, and 1000 U/ml leukemia inhibitory factor (LIF, Chemicon International) with a media change every day. Cells were then grown in aggregate cultures to form EBs in DMEM/F12 media supplemented with 10% knock-out serum replacement, 2.4% N2, 4500 mg/l Glucose, 2mM L-glutamine, and 0.1mM β-mercaptoethanol. EBs were formed for 4 d and then plated on 10ug/ml laminin-coated plates. After 24h of culture, the media was replaced by DMEM/F12, 3% Knockout serum, N2, Glucose, 1X Glutamine, 2-BME supplemented with 1% Insulin/Transferrin/Selenium with a media change every day. After 7 d, cells were dissociated by Accutase StemPro and plated on laminin-coated plates using a 1:1 ratio of Neurobasal media and DMEM/F12, N2, B27, 2mM L-glutamine, 0.1mM nonessential amino acids, 0.1 mM β-mercaptoethanol supplemented with 20 ng/ml bFGF (R&D Systems), 200 ng/ml SHH (R&D Systems) and 25ng/ml FGF8b (R&D Systems) with a media change every day. After 8 d, the culture medium was changed to Neurobasal/B27 medium supplemented with 0.5mM dbcAMP (Santa cruz), 0.2 mM ascorbic acid (Stem cell technologies), 20ng/ml BDNF (Petrotech) and 20ng/ml GDNF (Petrotech) with a media change every other day.

### Differentiation and culture of dopaminergic neurons from human induced-pluripotent stem cells (hiPSCs)

Primary fibroblasts derived from PD patients carrying the *GBA-N370S* mutation and a healthy control (below) were reprogrammed to pluripotency as described previously (Fernandes et al., 2016) and clones were selected, tested for mycoplasma and QCed according to established protocols (Lang et al., 2019). hiPSCs were differentiated toward dopaminergic fate as described by (Kriks et al., 2011) with small modifications (Beevers et al., 2017). Briefly, hiPSCs were patterned for 21 days with a growth factor cocktail to promote differentiation toward ventral midbrain neuronal progenitor cells for 11 days (10 mM SB431542, Tocris; 100 nM LDN193189, Sigma-Aldrich; 2 mM puromorphamine, Milipore; 100 ng/ml sonic hedgehog, Bio-Techne; 100 ng/ml fibroblast growth factor-8a, Bio-Techne and 3 mM CHIR99021), followed by 10 d of differentiation to dopaminergic neurons (20 ng/ml brain-derived neurotrophic factor, Peprotech; 20 ng/ml glial cell line-derived neurotrophic factor, Peprotech; 1 ng/ml transforming growth factor type β3, Peprotech; 0.5 mM dibutyryl cAMP, Sigma-Aldrich; 0.2 mM Ascorbic acid, Sigma-Aldrich and 10 mM DAPT, Abcam). Neurons were matured for a further 2 weeks (to day 35) for **α**-synuclein secretion, or for a further 5 weeks (to day 65) for analysis by SDS-PAGE and immunoblot. Neurons were then treated with DMSO, quercetin or 2-bromoplamitic acid for 3 d.

| Donor ID | Study ID | Genotype | Age / Gender | Characterisation |
| --- | --- | --- | --- | --- |
| SFC156-03 | Control | GBA wt /GBA wt | 75 Male | (Lang et al., 2019) |
| MK071 | PD 1 | GBA N370S /<br>GBA wt | 81 Female | (Fernandes et al., 2016) |
| MK088 | PD 2 | GBA N370S /<br>GBA wt | 46 Male | (Fernandes et al., 2016) |
| SFC871-03-09 | PD 3 | GBA N370S /<br>GBA wt | 70 Female | (Helle Bogetofte, 2021) |
| MK082 | PD 4 | GBA N370S /<br>GBA wt | 51 Male | (Lang et al., 2019) |

### Extracellular α-syn measurements

Commercial ELISA Kits as listed in Key Resources Table were used to quantify the extracellular α-syn in SH-SY5Y and mouse mDA neuronal culture. Conditioned media were collected and centrifuged at 1,000 g for 10 min. α−Syn in the recovered supernatant was measured using the protocol provided in the kit.

α-syn secretion by hiPSC-derived dopaminergic neurons was measured as described previously (Fernandes et al., 2016) (Fernandes et al., 2016) using an electro-chemiluminescent assay (Meso Scale Discovery, MD, USA, Cat# K151TGD-2) and a MESO QuickPlex SQ 120 instrument (Meso Scale Discovery) according to the manufacturer’s instructions. Briefly, culture media was collected 3d after drug treatment (differentiation day 38) and quantified relative to a standard curve. Data was normalized relative to the total protein content of the cells from which the conditioned media had been collected, as determined by BCA assay.

### Immunoblots

Cell lysate, cytosol or membrane samples, and extracellular vesicle samples were mixed with SDS sample loading buffer. Samples were heated at 95 °C for 5 min and separated on SDS-PAGE gels. Proteins were transferred to PVDF membranes (EMD Millipore, Darmstadt, Germany), blocked with 5% bovine serum albumin in TBST (20 mM Tris pH 7.4, 150 mM NaCl and 0.1% Tween-20) and incubated overnight with primary antibodies. For immunoblots from hiPSC dopamine neurons, samples in loading buffer were heated to 70°C for 10 minutes and blocking was carried out with 5% skimmed milk. For immunoblots of endogenous α-syn in SH-SY5Y cells, PVDF membranes were fixed with 0.4% paraformaldehyde (Electron Microscopy Science, Hatfield, PA) in TBST at room temperature for 30 min (Lee and Kamitani, 2011). Blots were then washed with TBST, followed by incubation with anti-rabbit or anti-mouse secondary antibodies (GE Healthcare Life Sciences, Pittsbugh, PA). Detection was performed with Supersignal^TM^ Chemiluminescent substrate (Thermo Fisher) and quantified with Fiji/ImageJ. Primary antibodies used in this study were listed in Key Resources Table. All antibodies used for immunoblots were diluted 1:1,000, except for 1:2,000 of mouse anti-Tubulin, 1:500 of rabbit anti-α-syn and of rabbit anti-tyrosine hydroxylase.

## Acknowledgements

We thank Dr. Michael Woodside and Dr. Thomas Südhof for sharing the plasmids, and Dr. Kalina Naidoo, Dr. Mootaz Salman and William McGuinness for preparation of iPSC samples. We also thank the staff at the UC Berkeley shared facilities, the cell culture facility, the DNA sequencing facility and Biological imaging facility. SW is supported as Associate of the HHMI. RS is an Investigator of the HHMI, a Senior fellow of the UC Berkeley Miller Institute of Science and the Scientific Director of Aligning Science Across Parkinson’s Disease.

## Additional information

### Funding

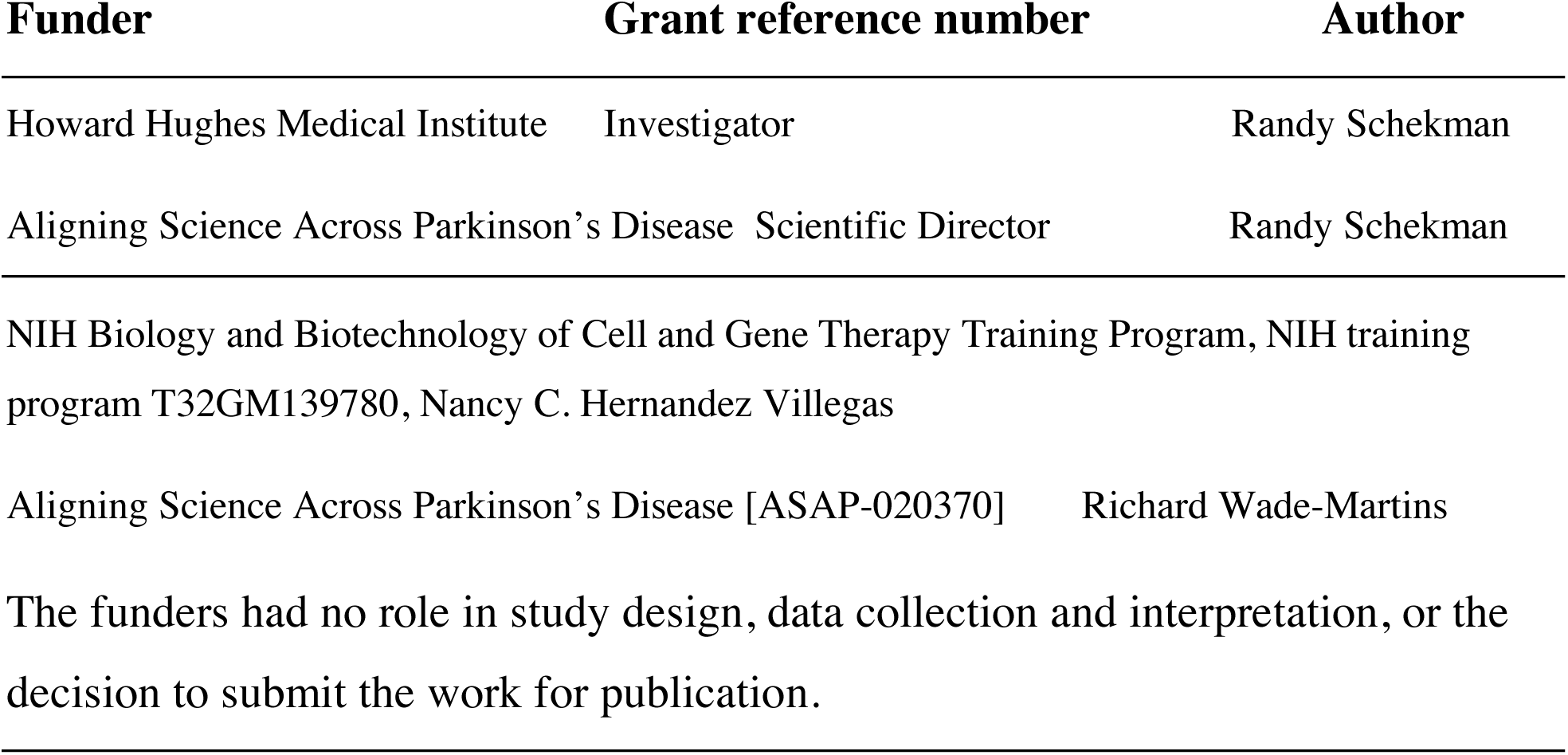

### Author Contributions

SW, Conception and design, Acquisition of data, Analysis and interpretation of data, Drafting or revising the article; DS, Conception and design, Acquisition of data, Analysis and interpretation of data, Drafting or revising the article; RS, Conception and design, Analysis and interpretation of data, Drafting or revising the article. NHV, Acquisition of data, analysis and interpretation of data, drafting or revising the article. IT-W, Acquisition of data, analysis and interpretation of data, drafting or revising the article. RW-M, Analysis and interpretation of data, drafting or revising the article.

---

**Figure 1 – source data**. Uncropped immunoblot and gel images corresponding to Figure 1.

**Figure 1 – figure supplement 1 – source data**. Uncropped immunoblot images corresponding to Figure 1 supplement 1.

**Figure 1 – figure supplement 2 – source data**. Uncropped immunoblot images corresponding to Figure 1 supplement 2.

**Figure 1 – figure supplement 3 – source data**. Uncropped immunoblot images corresponding to Figure 1 supplement 3.

**Figure 2 – source data**. Uncropped immunoblot corresponding to Figure 2.

**Figure 2 – figure supplement 1 – source data**. Uncropped immunoblot images corresponding to Figure 2 supplement 1.

**Figure 2 – figure supplement 2 – source data**. Uncropped immunoblot images corresponding to Figure 2 supplement 2.

**Figure 2 – figure supplement 3 – source data**. Uncropped gel images corresponding to Figure 2 supplement 3.

**Figure 3 – source data**. Uncropped immunoblot corresponding to Figure 3.

**Figure 3 – figure supplement 1 – source data**. Uncropped immunoblot corresponding to Figure 3 supplement 1.

**Figure 3 – figure supplement 2 – source data**. Uncropped immunoblot corresponding to Figure 3 supplement 2.

**Figure 4 – source data**. Uncropped immunoblot corresponding to Figure 4.

**Figure 5 – source data**. Uncropped immunoblot corresponding to Figure 5.

**Figure 5 – figure supplement 1– source data**. Uncropped immunoblot corresponding to Figure 5 supplement 1.

**Figure 5 – figure supplement 2– source data**. Uncropped immunoblot corresponding to Figure 5 supplement 2.

**Figure 6 – source data**. Uncropped immunoblot corresponding to Figure 6.

**Figure 6 – figure supplement 1 – source data**. Uncropped immunoblot corresponding to Figure 6 supplement 1.

**Figure 7 – source data**. Uncropped immunoblot corresponding to Figure 7.

**Figure 7 – figure supplement 1 – source data**. Uncropped immunoblot corresponding to Figure 7 supplement 1.

**Figure 7 – figure supplement 2 – source data**. Uncropped immunoblot corresponding to Figure 7 supplement 2.

**Figure 7 – figure supplement 3 – source data**. Uncropped immunoblot corresponding to Figure 7 supplement 3.

**Figure 7 – figure supplement 4 – source data**. Uncropped immunoblot corresponding to Figure 7 supplement 4.

**Figure 8 – source data**. Uncropped immunoblot corresponding to Figure 8.

**Figure 1 – figure supplement 1.**
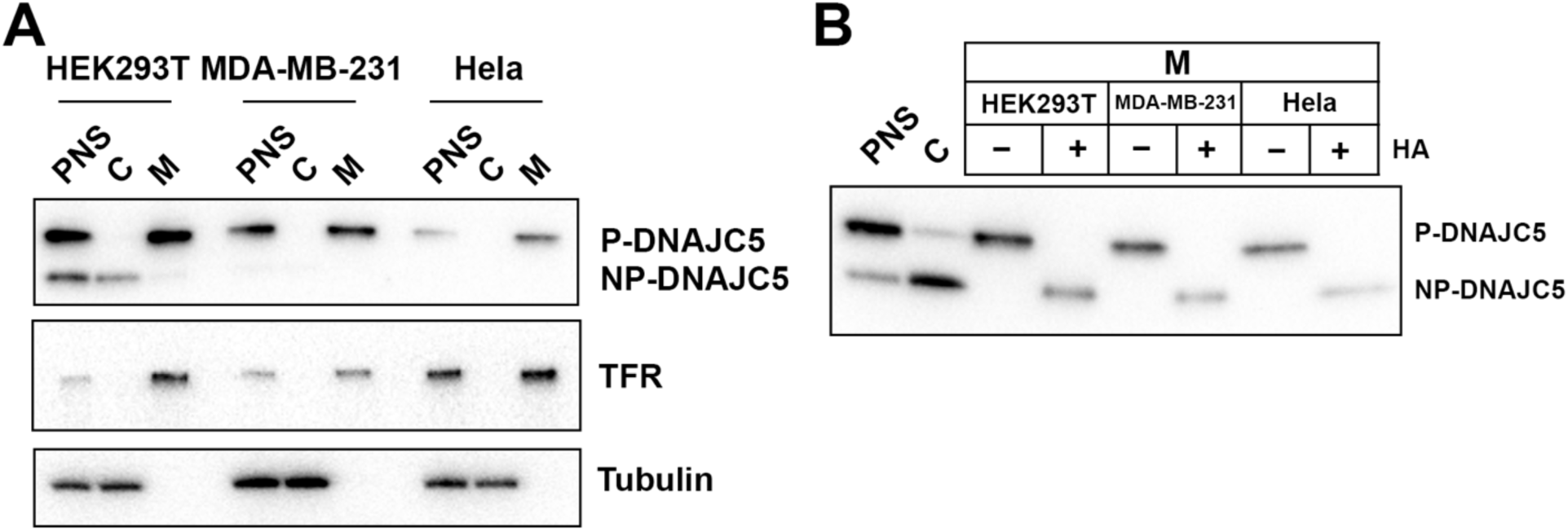
Validation of palmitoylation of DNAJC5 in various cell lines. (A) Membrane and cytosol fractionation of various cell lines (HEK293T, MDA-MB-231 and Hela) transfected with DNAJC5. The fractionation was performed as depicted in Figure 1B. PNS, post-nuclear supernatant. C, cytosol. M, membrane. Transferrin receptor (TFR) was used as a membrane marker. Tubulin was used as a cytosol marker. (B) In vitro depalmitoylation assay. Sedimented membranes (M) from different DNAJC5-transfected cell lines were collected and resuspended in 0.25M Tris pH 7.2 buffer or 0.25M Hydroxylamine (HA) pH 7.2 buffer. After overnight incubation at room temperature, the samples were examined by SDS-PAGE followed by anti-DNAJC5 immunoblot. PNS and C were used to compare the mobility of P-DNAJC5 and NP-DNAJC5, respectively.

**Figure 1 – figure supplement 2.**
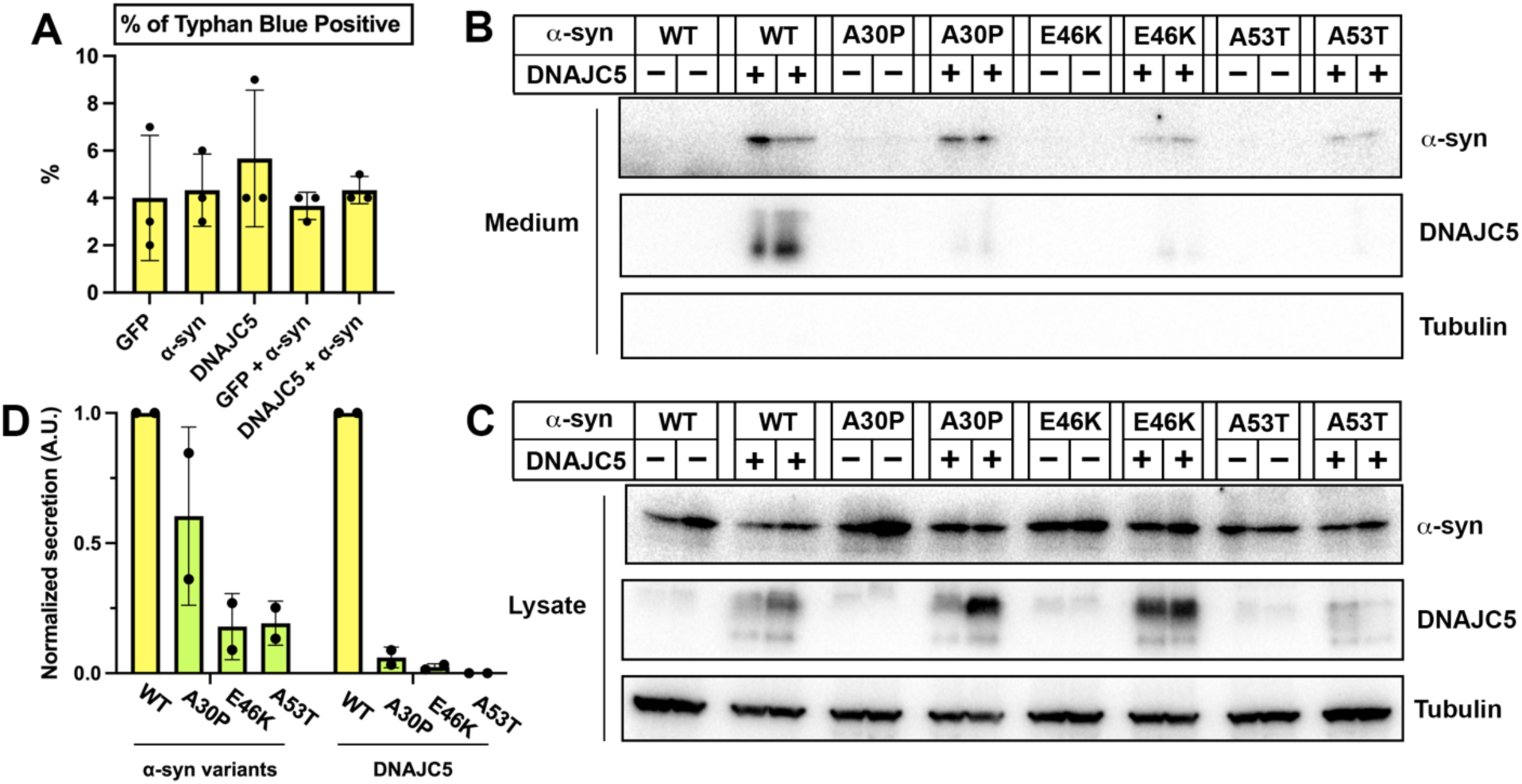
Secretion of α-syn variants. (A) Trypan blue cell vital staining after transfection with various constructs used in Figure 1. Ratios of trypan blue positive cells indicate the toxicity caused by transfection. Error bars represent standard deviations of 3 samples. (B) Secretion of α-syn variants into conditioned medium. Medium was collected, concentrated and evaluated by SDS-PAGE and immunoblot. (C) Expression of α-syn variants in HEK293T cells. HEK293T cells were co-transfected with Parkinson’s disease (PD)-causing α-syn mutant (A30P, E46K, A53T) and DNAJC5. (D) Quantification of normalized secretion (amount in medium divided by amount in lysate) of various α-syn variants and DNAJC5. The quantification was based on immunoblot in (B) and (C).

**Figure 1 – figure supplement 3.**
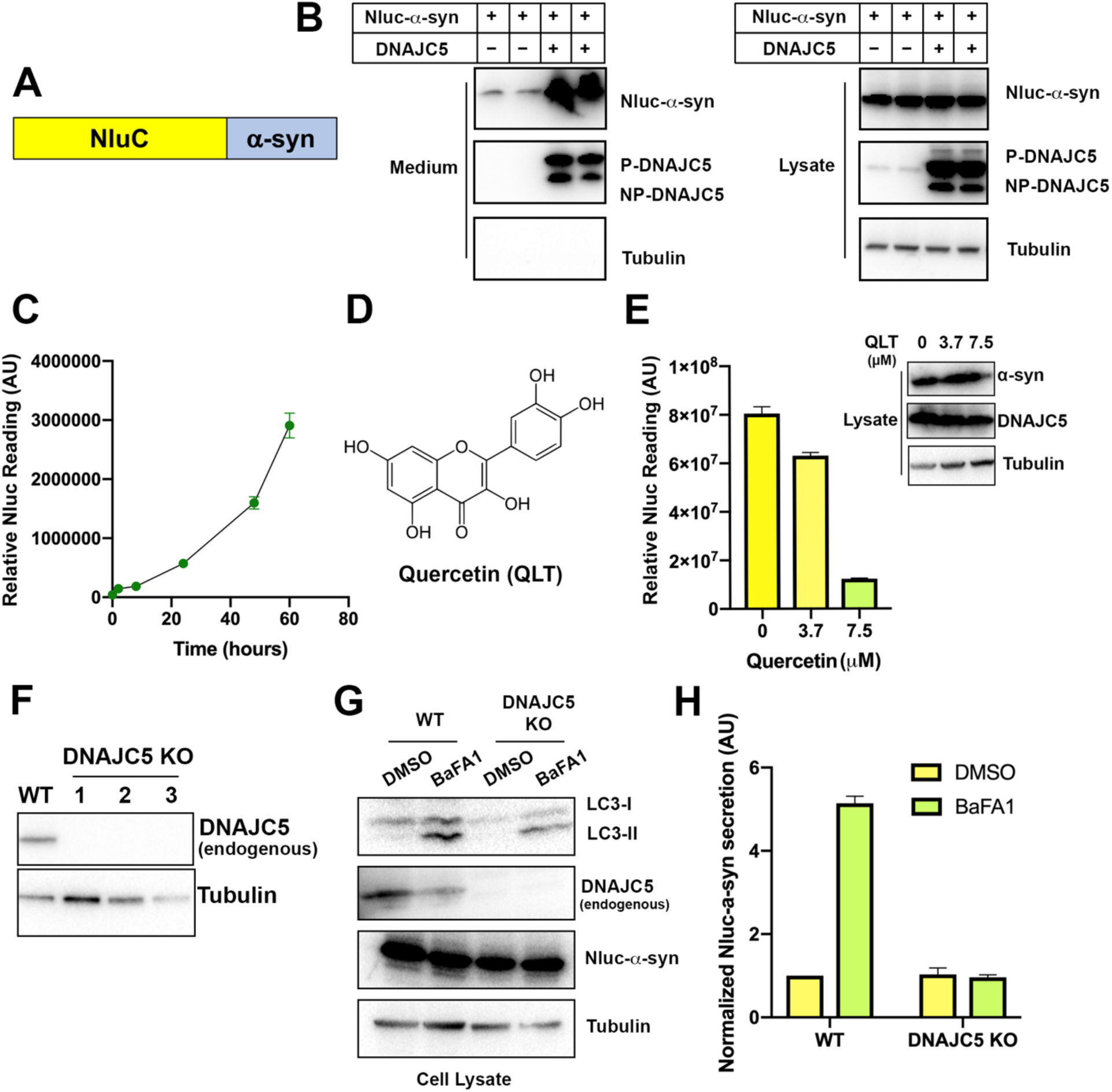
Secretion of α-syn is partially dependent on endogenous DNAJC5 in HEK293T cells. (A) Schematic diagram of nanoluciferase (Nluc)-fused α-syn. (B) Nluc-α-syn secretion was stimulated by DNAJC5 expression. Plasmids encoding Nluc-α-syn and DNAJC5 were co-transfected into HEK293T cells. Expression and secretion of proteins were detected with immunoblot 36 h after transfection. (C) Time-dependent of accumulation of Nluc-α-syn in the medium without DNAJC5 overexpression. After transfecting HEK293T cells with Nluc-α-syn alone, fractions of medium were collected at indicated time points. (D) Chemical structure of quercetin (QLT), a reported DNAJC5 inhibitor. (E) QLT inhibited endogenous Nluc-α-syn secretion in a concentration-dependent manner. AU, arbitrary unit. Secretion assay similar to (B) was performed with treatment of indicated concentration of QLT. Amounts of secreted proteins were quantified with nanoluciferase assay 36h after transfection. Immunoblot of α-syn and DNAJC5 in cell lysate after QLT treatment was shown on the right. Error bars represent standard deviations of three samples. (F) Validation of DNAJC5 knockout (KO) cell line generated by CRISPR. Wildtype (WT) HEK293T cell was used as a control. In all three single clones of DNAJC5 KO cell lines, DNAJC5 was not detectable by immunoblot. (G) Treatment of bafilomycin A1 (BaFA1) induced LC3 lipidation in cells. After 24h of 100 nM BaFA1 treatment, media were collected, and cells were lysed for evaluation with SDS-PAGE followed by immunoblot. The accumulation of the lipidated form of LC3 (LC3-II) was used to indicate the inhibition of autophagy and lysosomal degradation in cells. (H) Quantification of α-syn secretion from WT and DNAJC5 KO HEK293T cells after 24h treatment of BaFA1. The normalized secretion was calculated as nanoluciferase reading from media divided by the reading from cell lysate. Error bars represent standard deviations of three samples.

**Figure 2 – figure supplement 1.**
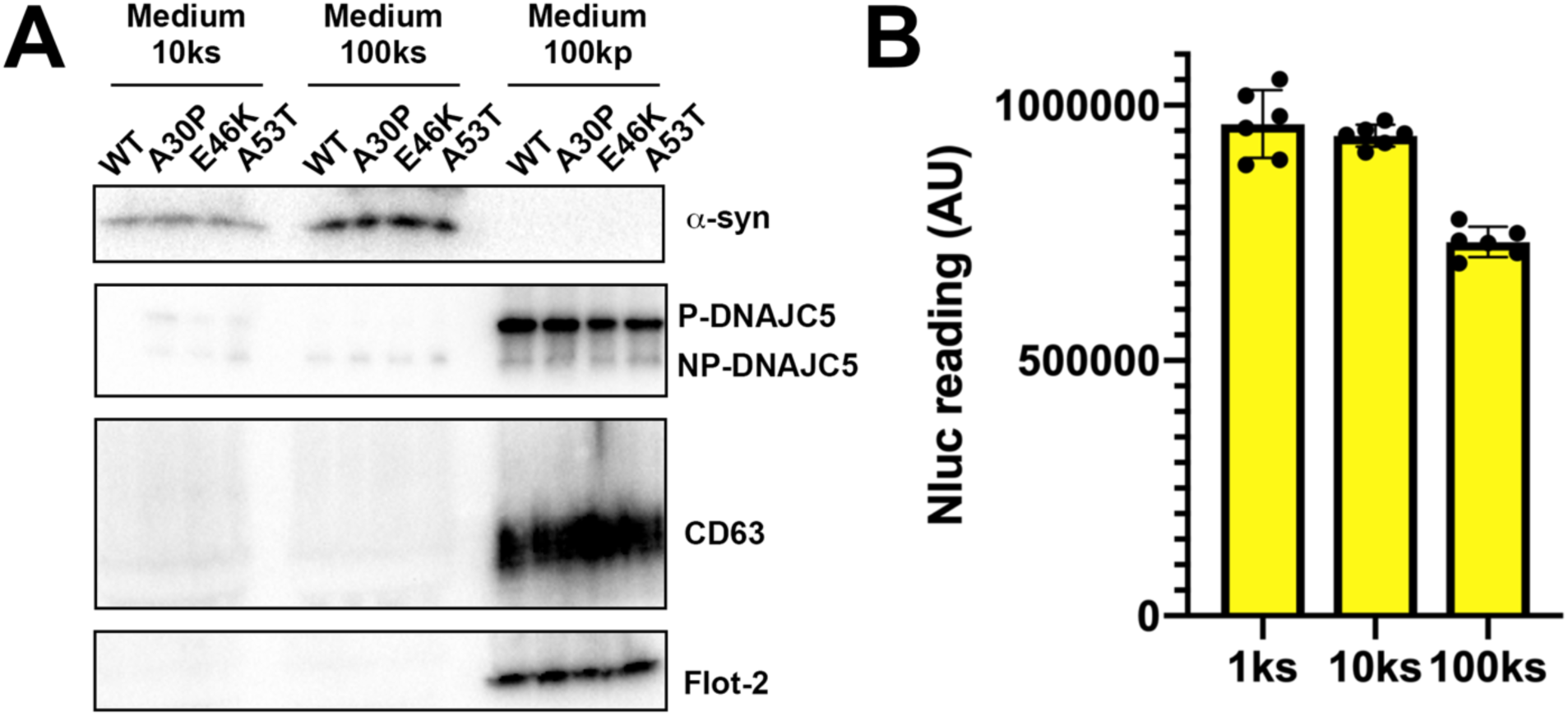
Solubility of secreted α-syn variants. (A) Medium fractionation of secreted α-syn PD mutants (A30P, E46K and A53T). After differential centrifugation of medium, supernatant (10ks and 100ks) and pellet fractions (100kp) were evaluated by immunoblot. (B) Medium fractionation of basal secreted Nluc-α-syn without DNAJC5 overexpression. Similar fractionation assay was performed in (A) with medium from HEK293T cells transfected with Nluc-α-syn alone. Secreted Nluc-α-syn was quantified with a nanoluciferase assay. AU, arbitrary unit.

**Figure 2- figure supplement 2.**
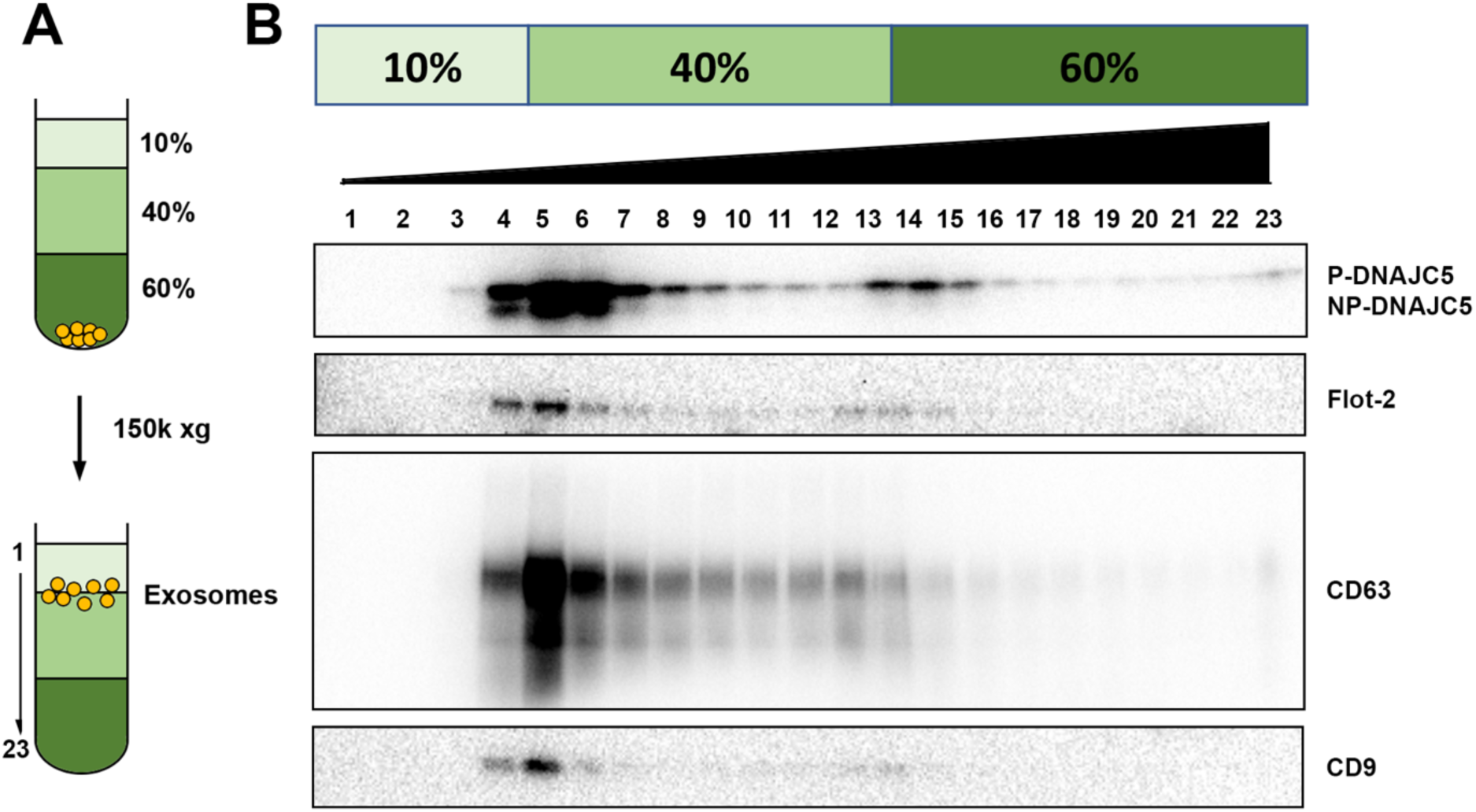
DNAJC5 enriched in buoyant EV fraction. (A) Schematic diagram of EV flotation protocol. Briefly, high-speed pellet fractions of growth medium were resuspended in 60% sucrose buffer and overlaid sequentially with 40% and 10% sucrose buffer. The tubes were centrifuged at 150,000 (150k)xg at 4°C for overnight. Buoyant EVs floated at the 10%/40% sucrose interface, separated from other insoluble materials. (B) Immunoblots across the sucrose step gradient. DNAJC5 as well as other classical exosome markers (Flot-2, CD63 and CD9) were highly enriched in fractions 4 to 6 at the 10%/40% interface.

**Figure 2 – figure supplement 3.**
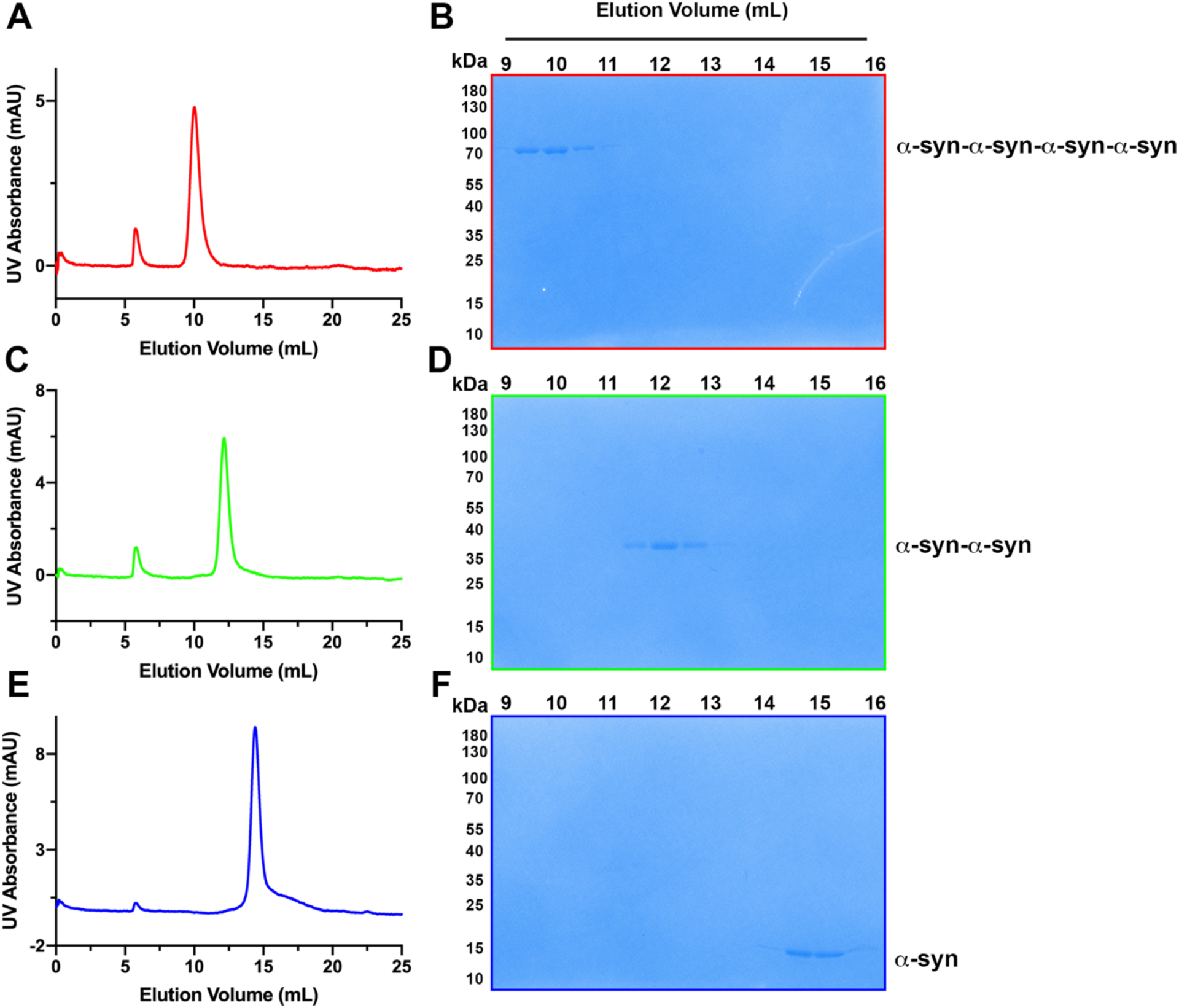
Assessment of tandem α-syn oligomers by gel filtration chromatography. (A) Chromatogram of purified tetrameric α-syn tandem oligomer (α-syn-α-syn- α-syn-α-syn). (B) Coomassie-blue stained SDS-PAGE of fractions from gel filtration of tetrameric α-syn tandem oligomer (α-syn-α-syn-α-syn-α-syn). (C) Chromatogram of purified dimeric α-syn tandem oligomer (α-syn-α-syn). (D) Coomassie-blue stained SDS-PAGE of fractions from gel filtration of dimeric α-syn tandem oligomer (α-syn-α-syn). (E) Chromatogram of purified WT α-syn (α-syn). (F) Coomassie-blue stained SDS-PAGE of fractions from gel filtration of WT α-syn (α-syn).

**Figure 3 – figure supplement 1.**
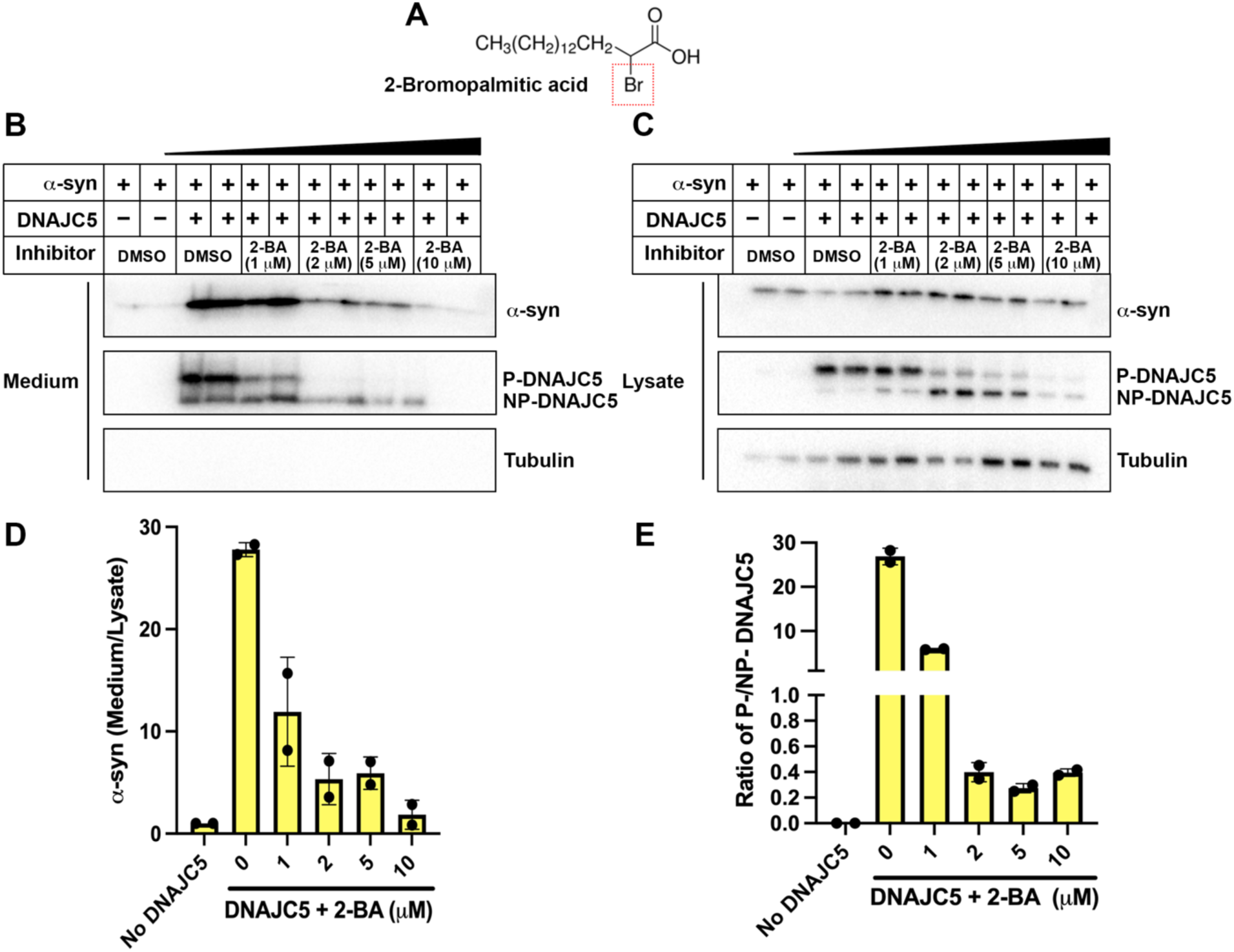
Dose-dependent inhibition of α-syn secretion by 2-bromopalmitic acid (2-BA). (A) Chemical structure of palmitoylation inhibitor 2-bromopalmitic acid (2-BA). The single bromo substituent at position 2 is highlighted by a red dashed square. (B) α-syn secretion in the medium was inhibited by 2-BA in a dose-dependent manner. Ten μm 2-BA was serially diluted by DMSO into 5 μm, 2μm, 1μm solution. HEK293T cells were first transfected with DNAJC5 and α-syn. After medium replacement, 2-BA of indicated concentration was added to cell culture. Media was collected after 36h and followed by sample preparation, SDS-PAGE and immunoblot. (C) With increasing concentration of 2-BA, P-DNAJC5 decreased and NP-DNAJC5 increased in HEK293T cells. (D) Quantification of normalized α-syn secretion upon increasing concentration of 2-BA. Quantification was based on immunoblot in (B) and (C). The α-syn secretion was calculated as the amount of α-syn in media divided by the amount in lysate. (E) Quantification of ratio of P-DNAJC5/NP-DNAJC5. The quantification was based on immunoblot in (C).

**Figure 3 – figure supplement 2.**
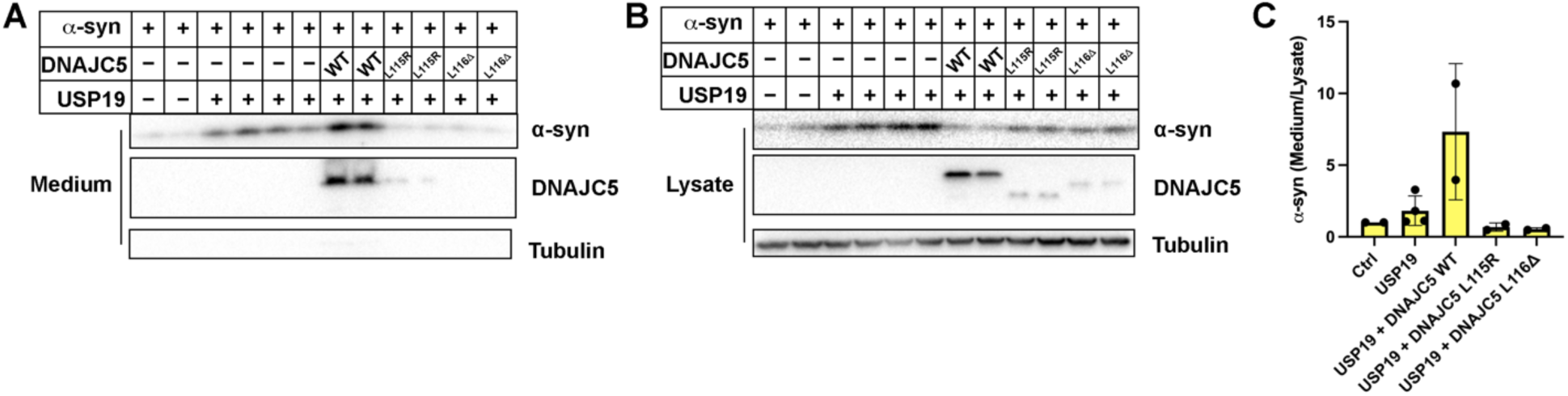
Blockage of USP19-induced α-syn secretion by DNAJC5 L115R or L116Δ mutant. (A) USP19 induced α-syn secretion in the medium, which was further enhanced by WT DNAJC5 and blocked by two DNAJC5 palmitoylation-deficient mutants (L115R and L116Δ). (B) Both mutations, L115R and L115Δ, inhibited DNAJC5 palmitoylation in HEK293T cells. (C) Quantification of normalized α-syn secretion in HEK293T cell transfected with indicated constructs. The quantification was based on immunoblot in (A) and (B). α-Syn secretion was calculated as the amount of α-syn in media divided by the amount in lysate.

**Figure 3 – figure supplement 3.**
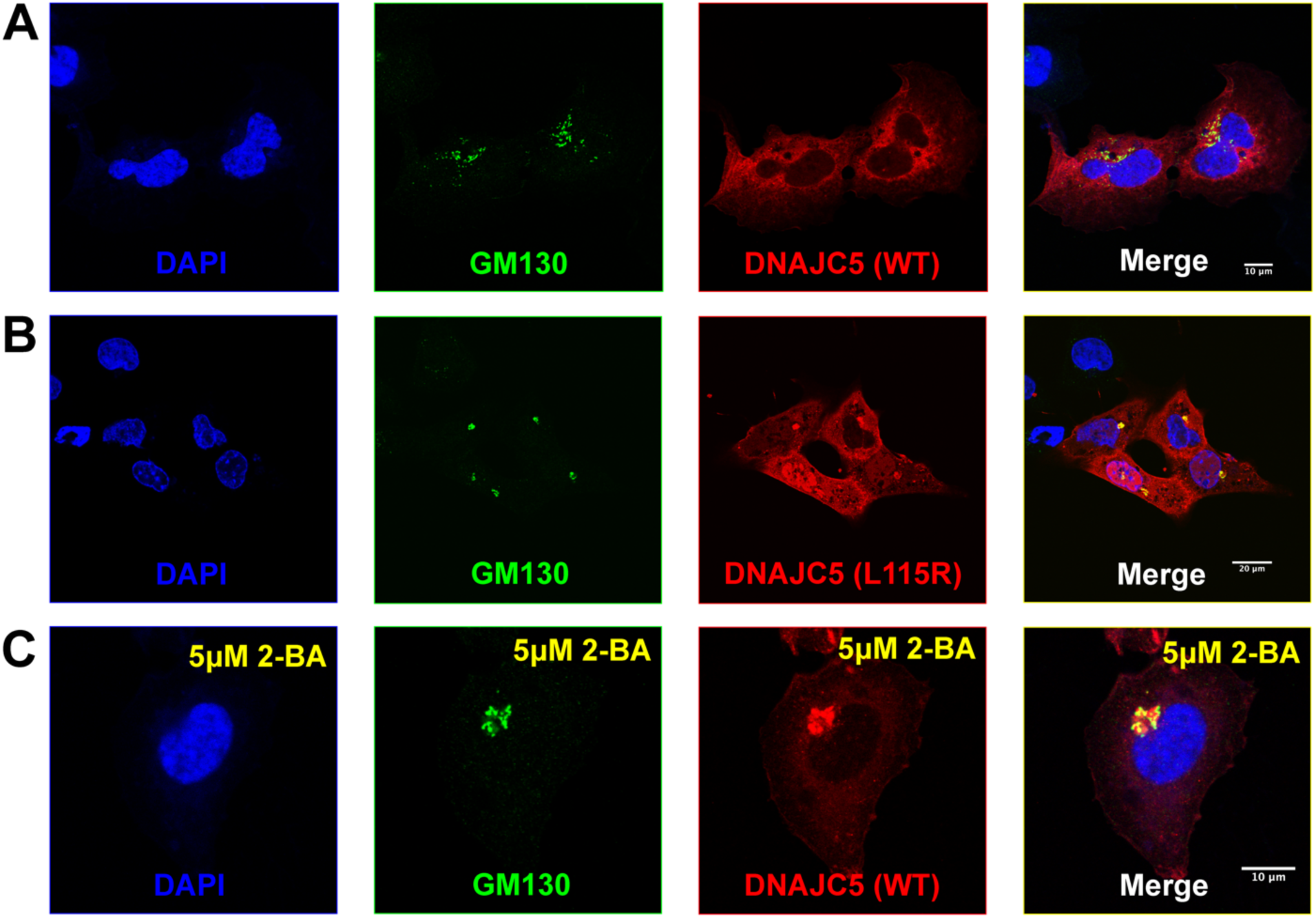
Golgi retention of DNAJC5 in secretion-deficient conditions. (B) Immunofluorescence (IF) images of U2OS cells transfected with wildtype (WT) DNAJC5-HaloTag. Before fixation, HaloTag TMR ligands were added to the cell culture to label DNAJC5. Golgi apparatus was visualized by IF using anti-GM130 antibody. Nuclei were stained with DAPI. Scale bar: 10 μm. (C) Immunofluorescence (IF) images of U2OS cells transfected with DNAJC5 (L115R)-HaloTag. Same staining procedure was performed as in (A). Scale bar: 20 μm. (D) Immunofluorescence (IF) images of U2OS cells transfected with wildtype (WT) DNAJC5-HaloTag and treated with 5 μM 2-BA. After transfection with WT DNAJC5-HaloTag, cells were incubated with 5μM 2-BA for 1d to block palmitoylation of DNAJC5. Same staining procedure was performed as in (A). Scale bar: 10 μm.

**Figure 4 – figure supplement 1.**
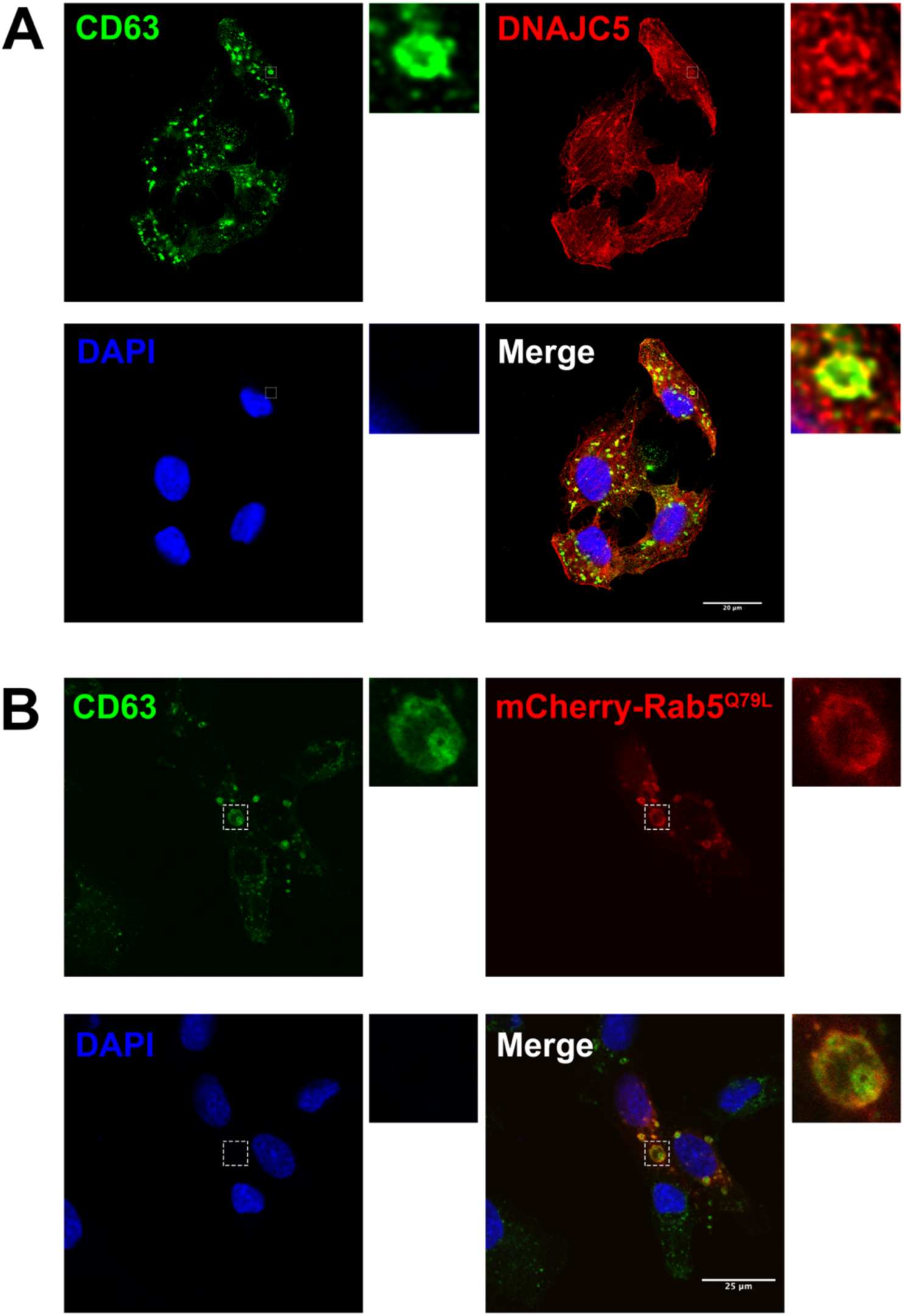
Immunofluorescence (IF) images of endogenous DNAJC5 and enlarged endosomes. (A) Co-localization between endogenous CD63 (red) and DNAJC5 (green). U2OS cells were cultured and fixed, incubated with corresponding antibodies for IF detection of endogenous CD63 and DNAJC5. Representative co-localized region is shown in magnified insets. Scale bar, 20 μm. (B) CD63 (green) was inside the enlarged endosomes labeled by peripheral mCherry-Rab5^Q79L^ (red). U2OS expressing mCherry-Rab5^Q79L^ were fixed, followed by IF using anti-CD63 antibody. Scale bar, 25 μm.

**Figure 4 – figure supplement 2.**
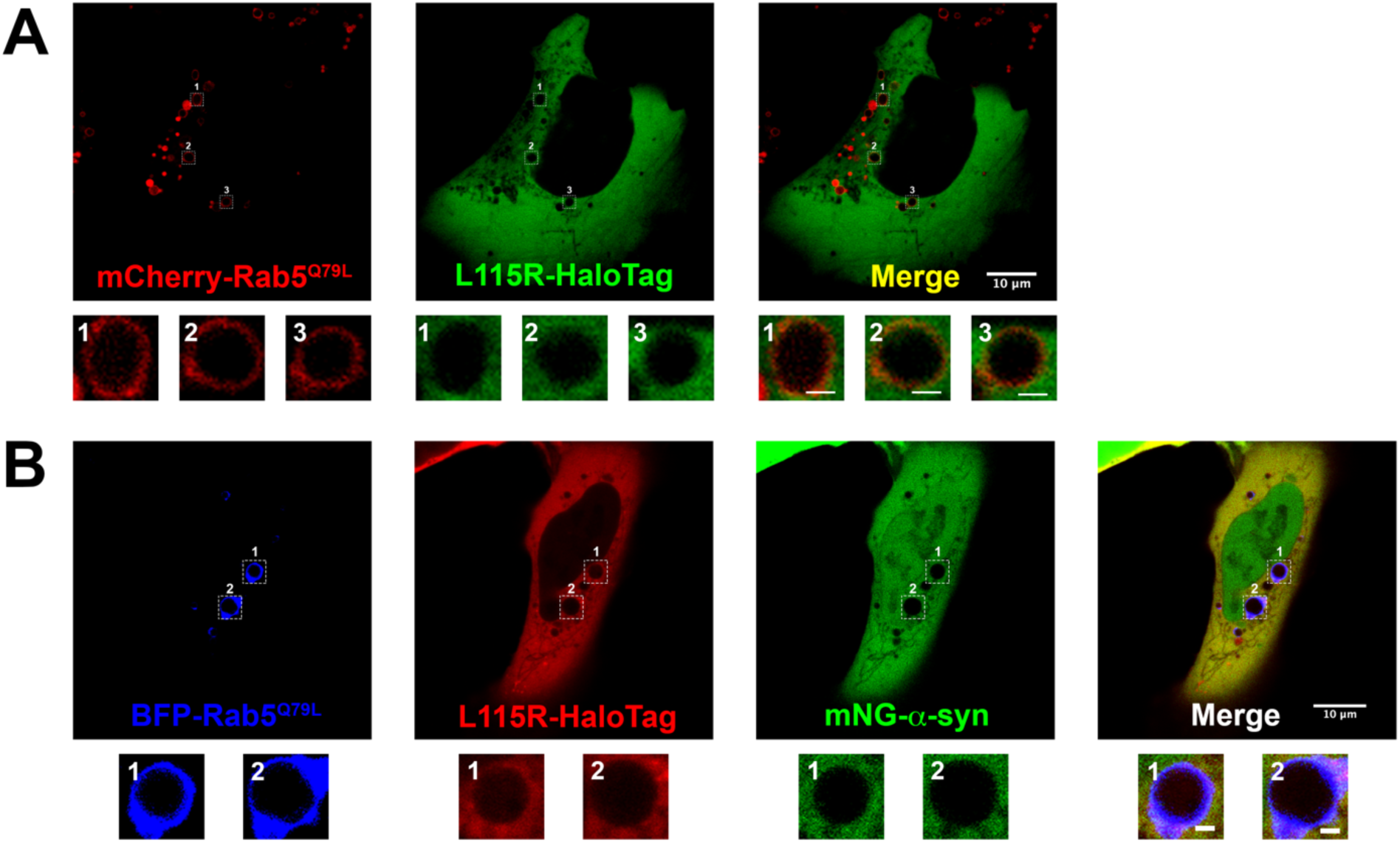
Live-cell images of U2OS cells expressing DNAJC5 L115R mutant and α-syn. (A) Live cell imaging of U2OS cells transfected with DNAJC5 (L115R)-HaloTag and mCherry-Rab5^Q79L^. DNAJC5 (L115R)-HaloTag (green) is diffuse in cytosol and outside of the enlarged endosomes labeled by peripheral mCherry-Rab5^Q79L^ (red). Scale bar, 10 μm in overviews and 1 μm in magnified insets. (B) Live cell imaging of U2OS cells transfected with mNeonGreen (mNG)-α-syn, DNAJC5 (L115R)-HaloTag and mCherry-Rab5^Q79L^. In the condition of co-expression with DNAJC5 (L115R)-HaloTag (red), no mNG-α-syn (green) was found inside of enlarged endosomes labeled by peripheral BFP-Rab5^Q79L^(blue). Scale bar, 10 μm in overviews and 1 μm in magnified insets.

**Figure 4 - supplement video.**
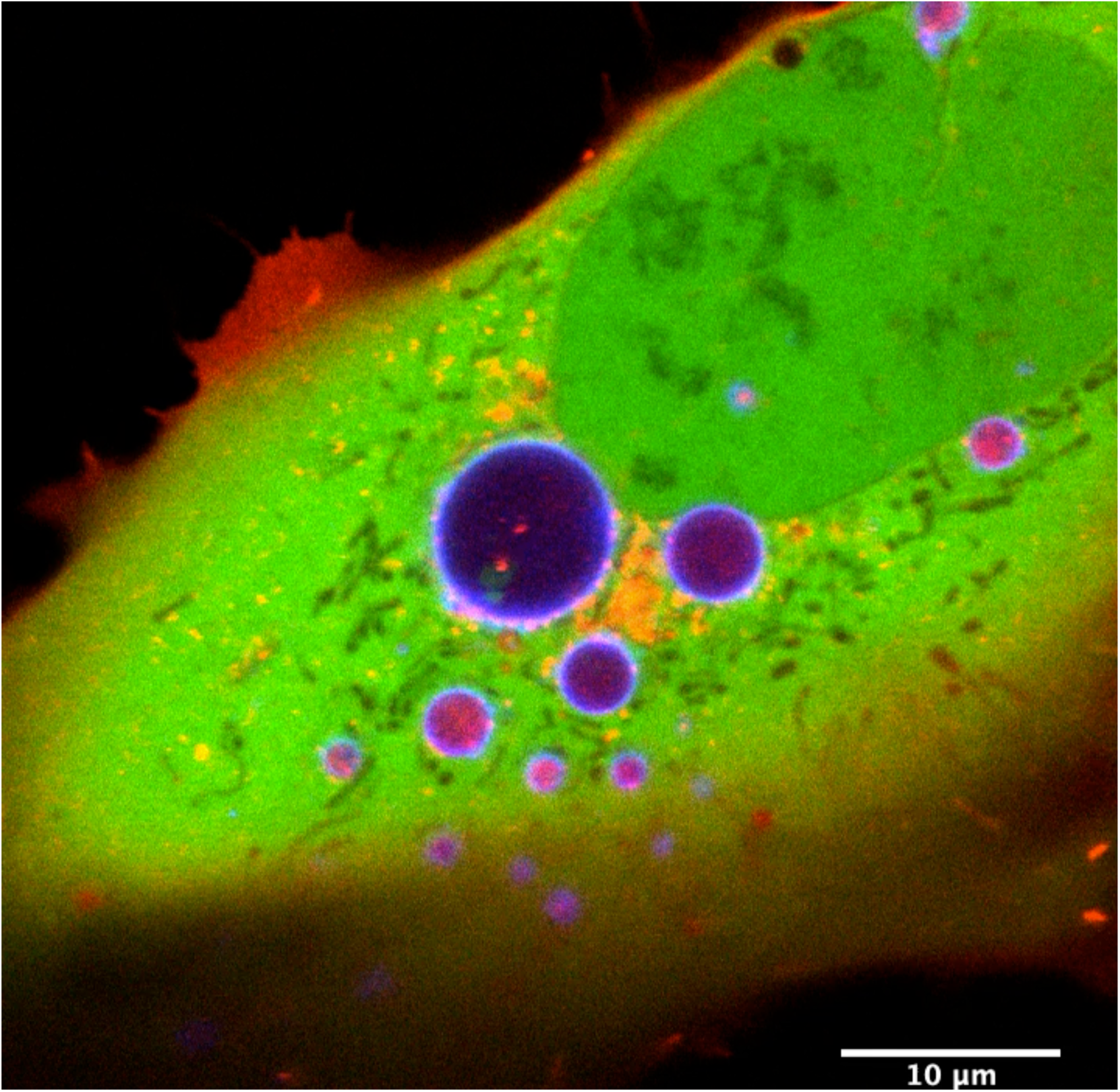
Time lapse of movement of internalized α-syn and DNAJC5 in the enlarged endosomes. DNAJC5-HaloTag (red) and mNG-α-syn (green) were co-expressed in U2OS cells carrying BFP-Rab5^Q79L^ (blue) mutant and imaged. Scale bar: 10 μm.

**Figure 5 – figure supplement 1.**
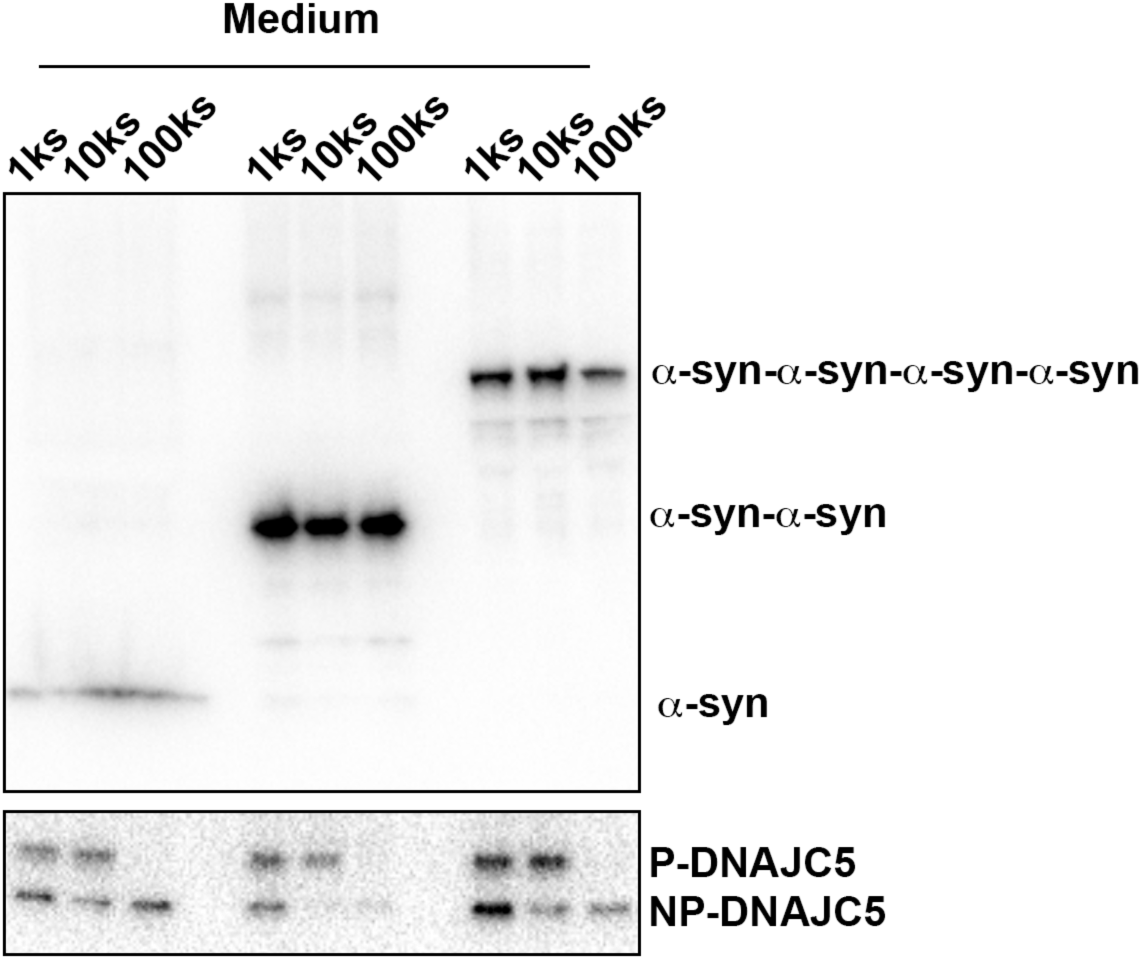
Medium fractionation of secreted tandem α-syn oligomers. Medium fractionation of secreted tandem α-syn oligomers. P-DNAJC5 was depleted in the supernatant after centrifugation at 100,000 (100k) xg. However, no significant decrease of tandem α-syn oligomers in the supernatant was observed.

**Figure 5 – figure supplement 2.**
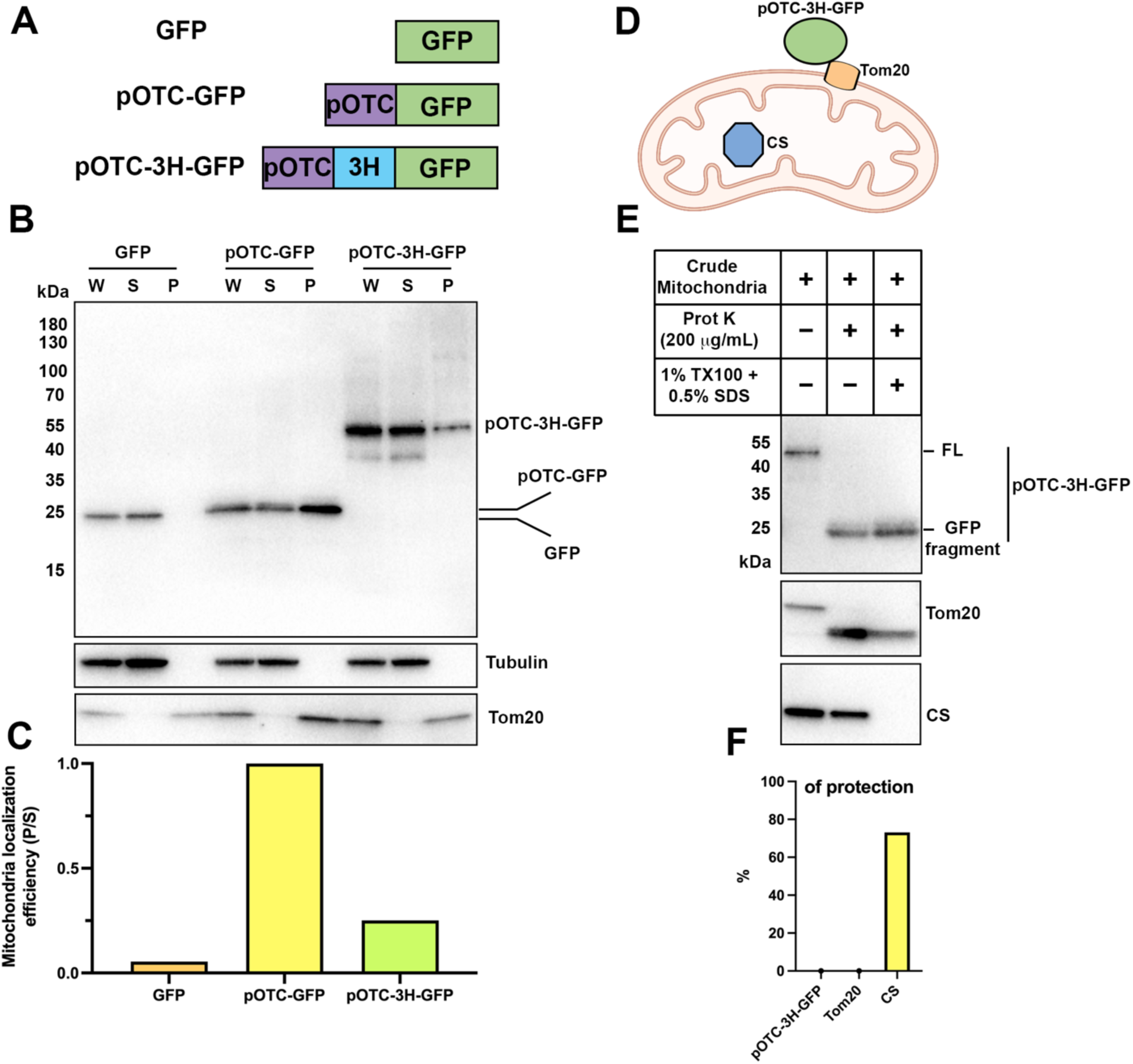
Thermostable three helix bundle blocked pOTC-mediated mitochondrial import of GFP. (A) Schematic diagrams of constructs used in mitochondrial import assay. GFP, GFP alone. pOTC-GFP, GFP with leader peptide from ornithine ranscarbamylase (OTC) fused at amino-terminus. pOTC-3H-GFP, the thermostable three helix bundle was inserted between the leader peptide pOTC and GFP. (B) Immunoblot analysis of proteins in crude mitochondria fraction. Whole cell lysate (W) was prepared by mixing homogenized cells with equal volume of 2.3M sucrose buffer and centrifugation at 1,200 g to remove large debris. Soluble fraction (S) and particulate fraction (P) were separated by centrifuging W fraction at 7,000 g for 10 min. Mitochondria were enriched in P fraction. Tom20, mitochondrial marker. Tubulin, soluble marker. (C) The mitochondrial localization efficiency was calculated by quantifying the protein amount in P fraction divided by the amount in S fraction. (D) Cartoon depicting relative localization of proteins inside or outside of mitochondria. Tom20, a mitochondrial outer membrane protein. Citrate synthase (CS), a mitochondrial matrix protein. The import of pOTC-3H-GFP is blocked by 3H. (E) Proteinase K protection assay of crude mitochondria. SDS (0.5%) was added in addition to 1% TX-100 to sensitize protease treatment of well-folded protein, e.g. CS. The protease accessibility of pOTC-3H-GFP was revealed by the production of GFP fragment and disappearance of the full-length (FL) band. (F) Quantification of percentage of protection of proteins in (E).

**Figure 6 – figure supplement 1.**
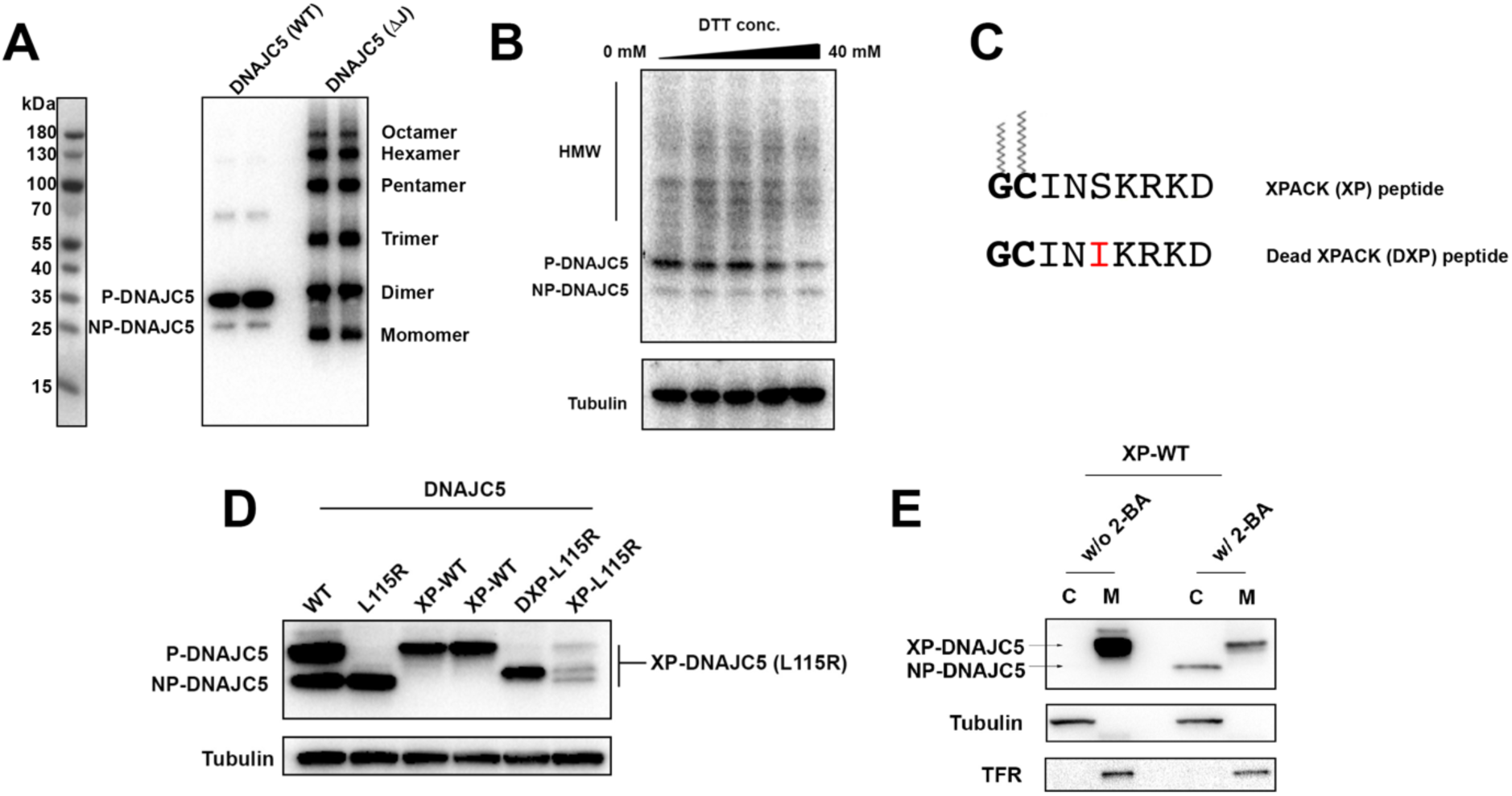
Characterization of HMW-DNAJC5 and XPACK fusion. (A) DNAJC5 (ΔJ) forms a series of SDS-resistant oligomers. HEK293T cells transfected with WT DNAJC5 or DNAJC5 (ΔJ) were lysed and evaluated by immunoblot using anti-DNAJC5 antibody. (B) HMW-DNAJC5 is not formed by non-specific disulfide bonds. HEK293T cells transfected with DNAJC5 were lysed for SDS-PAGE followed by immunoblot. The loading samples for SDS-PAGE were prepared with increasing amount of DTT up to 40 mM. (C) Schematic diagram of XPACK. XPACK is myristoylated on the first glycine (G) and palmitoylated on the second cystine (C). Replacement of Serine (S) at position 5 with a hydrophobic isoleucine (I) abolishes normal lipidation of XPACK. (D) Assessment of different XPACK fusion constructs. Cell lysate containing different DNAJC5 constructs were evaluated by anti-DNAJC5 immunoblot. (E) Membrane-association of XP-DNAJC5 dependent on palmitoylation. HEK293T cells transfected with XP-DNAJC5 were treated with DMSO or 10 μM 2-BA. After cell culture for 24 h, cellular fractionation was performed with homogenized cells. Distribution of XP-DNAJC5 in cytosol (C) and membrane (M) fractions were evaluated with immunoblot.

**Figure 6 – figure supplement 2.**
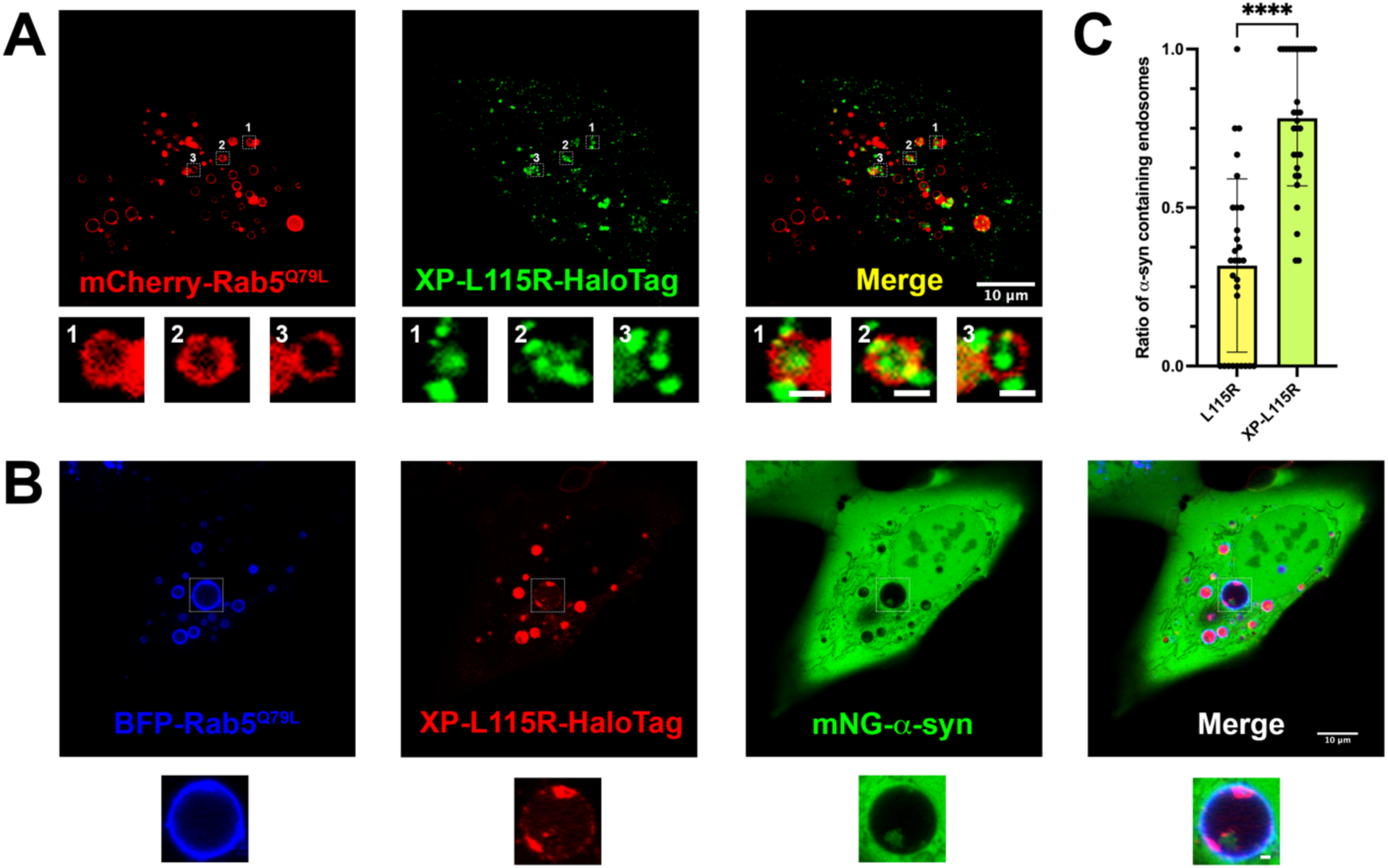
Live-cell images of U2OS cells expressing XP-DNAJC5 L115R mutant and α-syn. (A) Internalization of punctate XP-DNAJC5 (green) into enlarged endosomes labeled by mCherry-Rab5^Q79L^ (red). U2OS cells were transfected with XP-DNAJC5 and mCherry-Rab5^Q79L^. Live cell imaging was performed 24 h after transfection. Representative enlarged endosomes with XP-DNAJC5 are shown in magnified inset. Scale bar: 10 μm in overviews and 1 μm in magnified insets. (B) α-syn enters into enlarged endosomes in the presence of XP-DNAJC5. XP-DNAJC5-HaloTag (red) and mNG-α-syn (green) were co-expressed in U2OS cells carrying BFP-Rab5^Q79L^ (blue) mutant and imaged. Representative enlarged endosome with both XP-DNAJC5-HaloTag and mNG-α-syn inside is shown in magnified inset. Scale bar: 10 μm in overviews and 1 μm in magnified insets. (C) Quantification of the ratio of α-syn containing endosomes in cells co-transfect with DNAJC5 L115R or cells co-transfected with DNAJC5 XP-L115R. More than 100 enlarged endosomes were counted in each group. Error bars represent standard deviations. ****p value<0.0001, two-tailed t test.

**Figure 7 – figure supplement 1.**
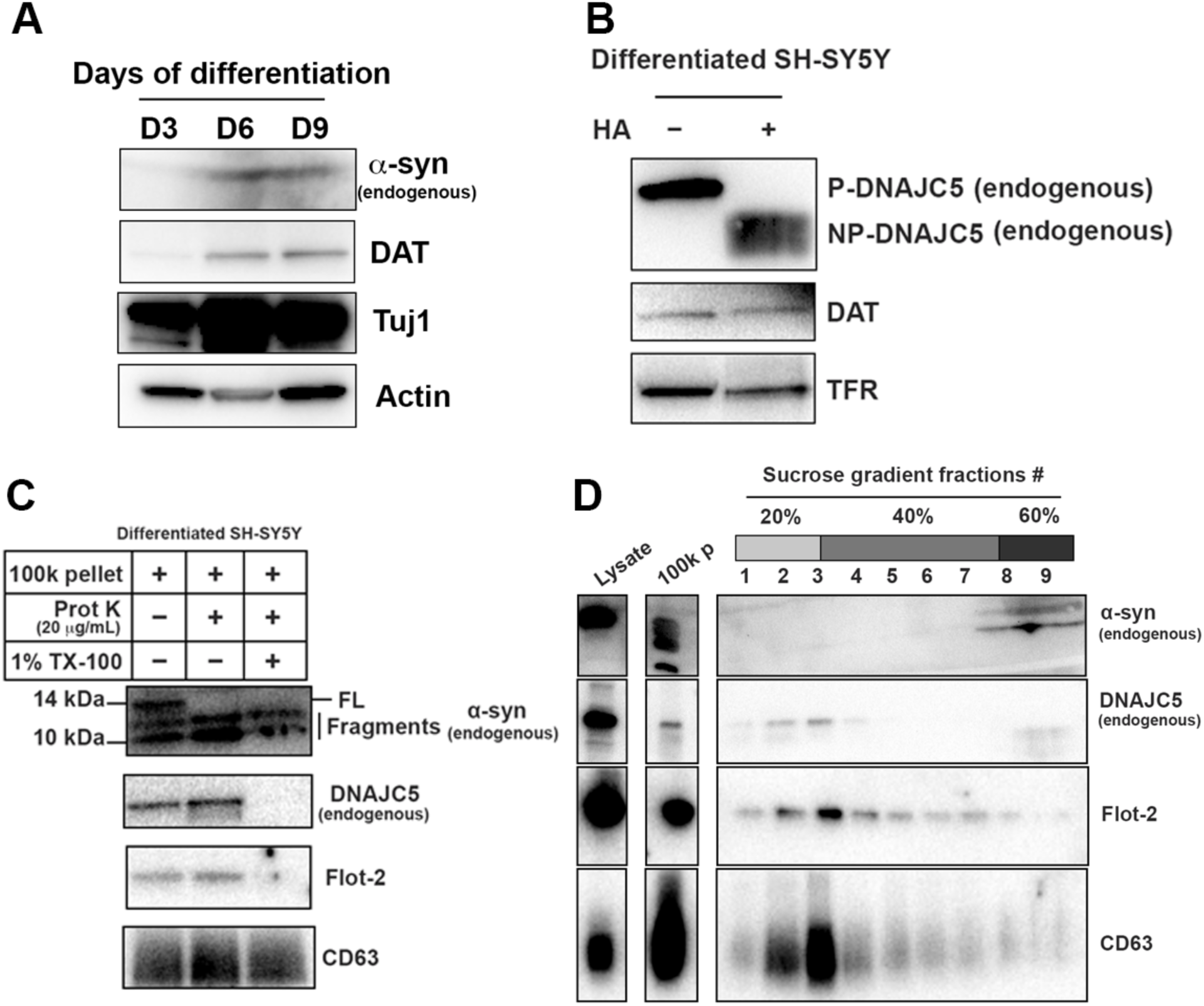
Basal α-syn secreted as a soluble form from differentiated SH-SY5Y. (A) Differentiation of SH-SY5Y was initiated by lowering the serum concentration to 1% FBS and addition of 10uM retinoic acid (RA). Media were replaced every three days and the cells were harvested at indicated time to examine the expression of neuronal marker. DAT, dopamine transporter. Tuj1, neuron-specific class III β-tubulin. (B) In vitro depalmitoylation assay of endogenous DNAJC5 in differentiated SH-SY5Y cells. The depalmitoylation assay was performed as in Fig 1 **supplement 1B** using membrane (M) fraction from SH-SY5Y cells. HA, hydroxylamine. (C) Proteinase K protection assay of 100,000 (100k) pellet fraction from the centrifuged media of differentiated SH-SY5Y culture. Flot-2 and CD63 were used as exosome markers. (D) Sedimented α-syn was not buoyant. 100k pellet from (C) was mixed with 60% sucrose in PBS and layered with 40% and 20% sucrose in PBS sequentially. After centrifuged at 150,000xg for 16 h, fractions were collected from top to bottom.

**Figure 7 – figure supplement 2.**
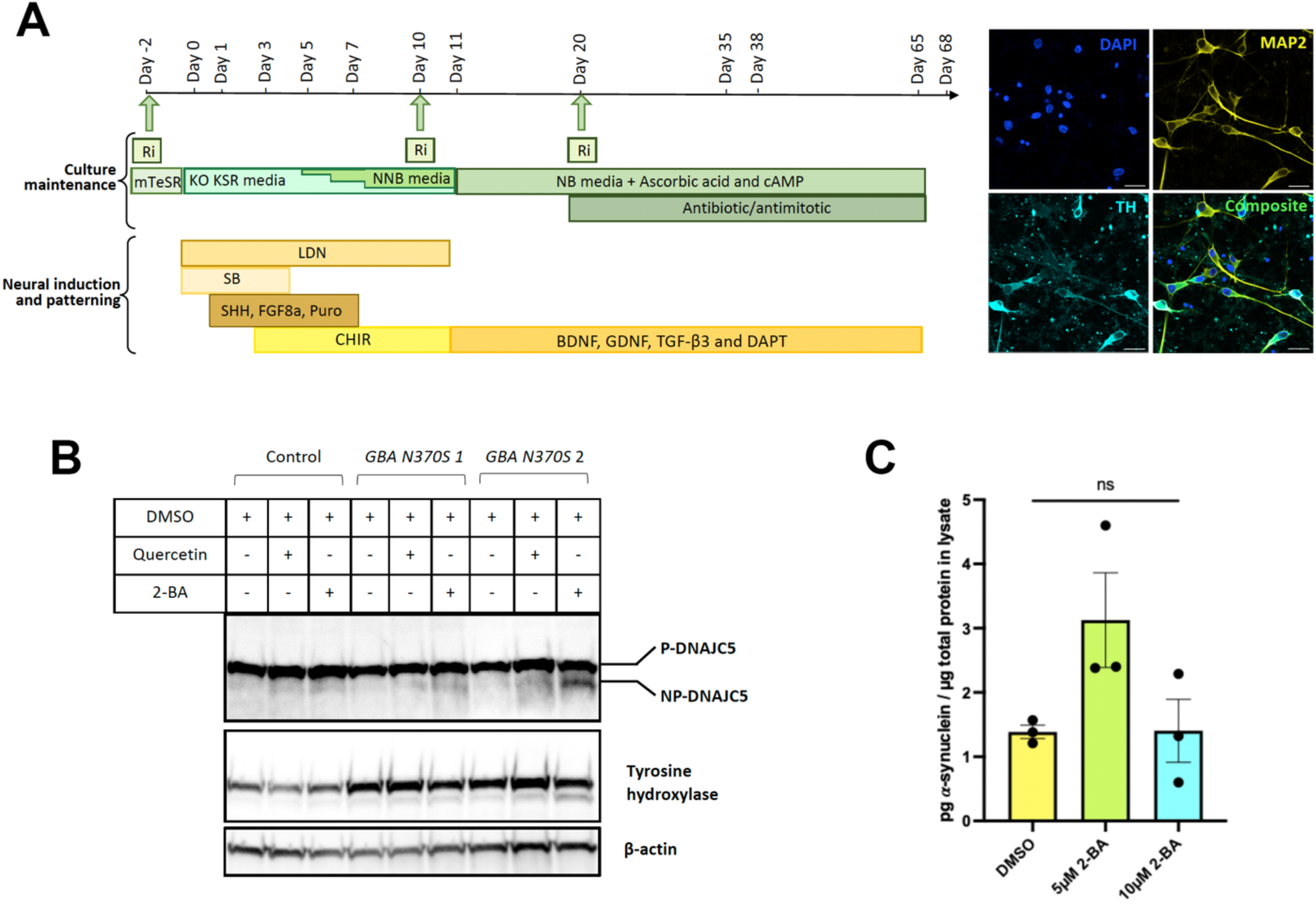
(A) Schematic of the differentiation protocol used to generate hiPSC-derived dopamine neurons, including a patterning phase to generate neural progenitor cells followed by differentiation into mature neurons. Green arrows represent points of replating when cells are supplemented with ROCK inhibitor (Ri) to increase survival. Puro = puromorphamine, SHH = sonic hedgehog, FGF8a = fibroblast growth factor 8a, BDNF = brain-derived neurotrophic factor, GDNF = glial cell line-derived neurotrophic factor, TGF-β3 = transforming growth factor beta 3. (see Kriks et al, 2011; Beevers et al 2017; Lang et al 2019 for full details). Neurons are mature and harvested between 35-68 d for analysis. Immunocytochemical fluorescent images of mature neurons at Day 50 stained for neuronal marker microtubule-associated protein (MAP2), the dopaminergic marker tyrosine hydroxylase (TH), and DAPI. Scale bar = 20 µm. (B) DNAJC5 is palmitoylated in iPSC-derived dopamine neurons. Immunoblot analysis of DNAJC5 palmitoylation in hiPSC-derived dopamine neurons treated with DMSO, 7.5 μM Quercetin or 10 μM 2-BA on Day 65 and harvested on Day 68. Tyrosine hydroxylase was used as a marker of dopaminergic identity. (C) Partial depalmitoylation of DNAJC5 by 2-BA in iPSC-derived dopamine neurons does not reduce α-syn secretion. hiPSC-derived dopamine neurons carrying the *GBA-N370S* mutation were treated with palmitoylation inhibitor 2-BA (5 μM or 10 μM) at Day 35. Culture media samples were harvested after 3d treatment at Day 38 and α-syn levels in the media were analyzed by electro-chemiluminescent immunoassay. Data points represent individual cell lines derived from different donors and are normalized to total protein in the corresponding cell lysates. One-way ANOVA shows no effect of 2-BA on α-syn secretion.

**Figure 7 – figure supplement 3.**
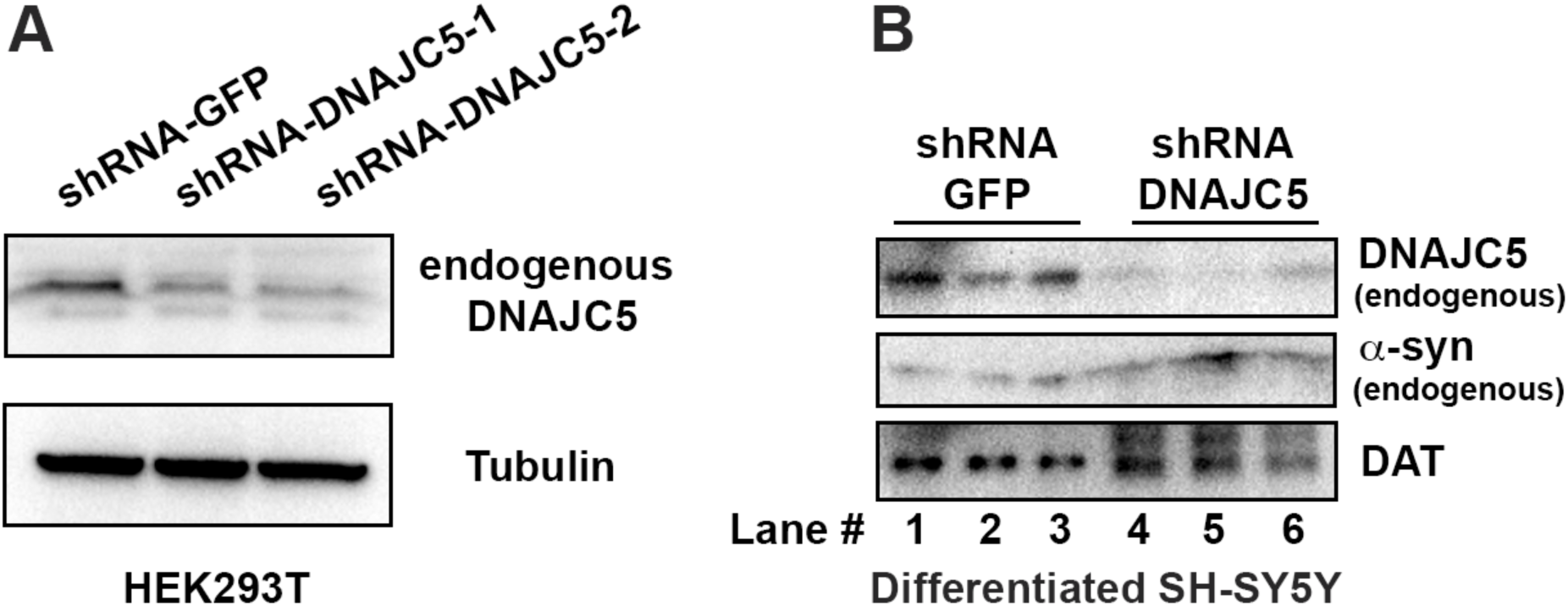
shRNA-mediated DNAJC5 knockdown in differentiated SH-SY5Y cells. (A) Examination of knockdown efficiency by shRNA targeting DNAJC5 in HEK293T cells. (B) Endogenous DNAJC5 expression decreased in SH-SY5Y cells transduced with shRNA targeting DNAJC5.

**Figure 7 – figure supplement 4.**
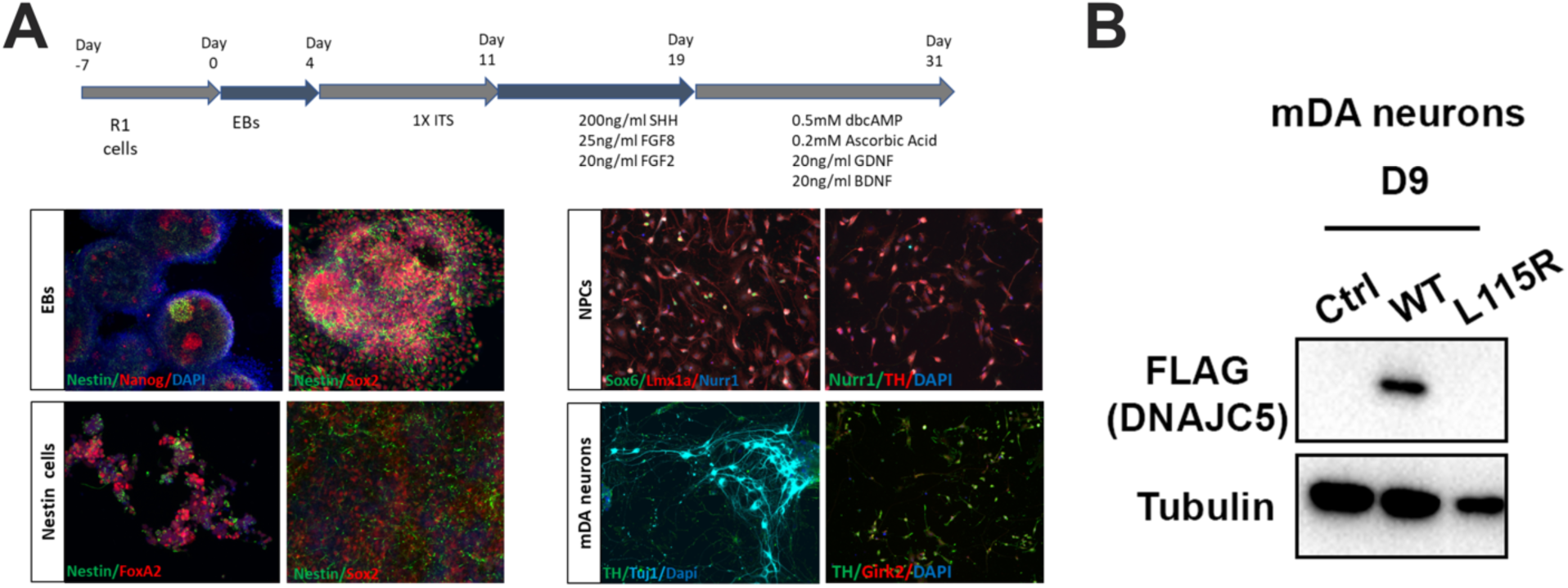
Differentiation of mouse embryonic stem cells (mESCs) and expression of human DNAJC5. (A) Schematic overview of protocol used for differentiation of mESCs into middle-brain dopaminergic (mDA) neuronal cultures. Immunocytochemical staining using stem cell markers (Nanog, Sox2), neuronal precursor marker (Nestin), mDA markers [FoxA2, Nurr1, Lmx1a, Sox6 (selective marker of substantia nigra pars compacta lineage), TH (tyrosine hydroxylase), Girk2 (G-protein-regulated inward-rectifier potassium channel 2, expressed in DA neurons)], neuronal marker (Tuj1) and nuclear marker (dapi). (B) Expression of human DNAJC5 with C-terminal FLAG tag in mDA neurons.

